# Transcriptomic diversity of amygdalar subdivisions across humans and nonhuman primates

**DOI:** 10.1101/2024.10.18.618721

**Authors:** Michael S. Totty, Rita Cervera Juanes, Svitlana V. Bach, Lamya Ben Ameur, Madeline R. Valentine, Evan Simons, McKenna Romac, Hoa Trinh, Krystal Henderson, Ishbel Del Rosario, Madhavi Tippani, Ryan A. Miller, Joel E. Kleinman, Stephanie Cerceo Page, Arpiar Saunders, Thomas M. Hyde, Keri Martinowich, Stephanie C. Hicks, Vincent D. Costa

**Affiliations:** Department of Biostatistics, Johns Hopkins Bloomberg School of Public Health, Baltimore, MD, USA; Lieber Institute for Brain Development, Johns Hopkins Medical Campus, Baltimore, MD, USA; Department of Translational Neuroscience, Center for Precision Medicine, Wake Forest University School of Medicine, Winston-Salem, NC, USA; Vollum Institute, Oregon Health and Science University, Portland, OR, USA; Division of Neuroscience, Oregon National Primate Research Center, Beaverton, OR, USA; Division of Developmental and Cognitive Neuroscience, Emory National Primate Research Center, Atlanta, GA, USA; Department of Neurology, Johns Hopkins School of Medicine, Baltimore, MD, USA; Department of Psychiatry and Behavioral Sciences, Johns Hopkins School of Medicine, Baltimore, MD, USA; The Solomon H. Snyder Department of Neuroscience, Johns Hopkins School of Medicine, Baltimore, MD, USA; Johns Hopkins Kavli Neuroscience Discovery Institute, Baltimore, MD, USA; Department of Biomedical Engineering, Johns Hopkins University, Baltimore, MD, USA; Center for Computational Biology, Johns Hopkins University, Baltimore, MD, USA; Malone Center for Engineering in Healthcare, Johns Hopkins University, Baltimore, MD, USA; Department of Psychiatry and Behavioral Sciences, Emory University, Atlanta, GA 30329, USA

## Abstract

The amygdaloid complex mediates learning, memory, and emotions. Understanding the cellular and anatomical features that are specialized in the amygdala of primates versus other vertebrates requires a systematic, anatomically-resolved molecular analysis of constituent cell populations. We analyzed five nuclear subdivisions of the primate amygdala with single-nucleus RNA sequencing in macaques, baboons, and humans to examine gene expression profiles for excitatory and inhibitory neurons and confirmed our results with single-molecule FISH analysis. We identified distinct subtypes of *FOXP2^+^* interneurons in the intercalated cell masses and protein-kinase C-δ interneurons in the central nucleus. We also establish that glutamatergic, pyramidal-like neurons are transcriptionally specialized within the basal, lateral, or accessory basal nuclei. Understanding the molecular heterogeneity of anatomically-resolved amygdalar neuron types provides a cellular framework for improving existing models of how amygdalar neural circuits contribute to cognition and mental health in humans by using nonhuman primates as a translational bridge.

## INTRODUCTION

The amygdala is a small structure in the temporal lobe containing multiple nuclear subdivisions. The majority of amygdala neurons are located in the lateral, basal, accessory basal, intercalated and central nuclei. These five nuclei define the primary input and output pathways for amygdala-dependent information processing. In rodents, individual amygdala nuclei are implicated in learning ^1^, decision-making ^2^ and social behaviors ^3^. Recent uses of molecular, genetic, and transcriptomic tools have revealed how unique cell types contribute to function in these individual nuclei ^4–8^.

The amygdala similarly mediates cognitive and emotional behaviors in nonhuman primates and humans ^9–13^. However, the more complex spatial arrangement of the primate amygdala has made it challenging to examine the functional effects of perturbing specific nuclei ^14^, or cell types in these nuclei. Targeting specific nuclei with neuronal tracers in combination with morphological analyses in the nonhuman primate, has provided clear descriptions of the connectivity of glutamatergic and GABAergic neurons across the major amygdalar nuclei ^15–17^, but the molecular composition of these cells within and across nuclear subdivisions is still not clear. Using a similar approach in nonhuman primates that incorporates transcriptomics would be invaluable for making inferences about the connectivity of molecularly-defined cell types in the human amygdala. However, recent efforts to profile neuron types in marmosets ^18^, macaques ^19,20^, or humans ^21,22^ lacked detailed anatomical context or utilized marker genes derived from rodent atlases for spatial annotations ^20^. While emerging spatial transcriptomic approaches can provide anatomical information, there remain computational and logistical challenges in using them to map gene expression in spatially complex structures like the amygdala ^23,24^.

This study constructed a spatially-resolved, transcriptomic atlas of the primate amygdala using single-nuclei RNA sequencing (snRNAseq) and single-molecule fluorescence *in situ* hybridization (smFISH). We systematically sampled tissue using cytoarchitectonic landmarks from the lateral, basal, accessory basal, and central nucleus in rhesus macaques and baboons (Fig. 1). Because *FOXP2* is a primate-conserved marker of interneurons in the intercalated cell masses (ITC) ^25^, which form a diffuse net between the central, accessory basal, basal, and lateral nuclei in macaques ^26^, our strategy allowed us to identify and profile this fifth subdivision. We used the macaque and baboon transcriptomic data to make inferences about the heterogeneity of glutamatergic and GABAergic neurons across nuclear subdivisions (Fig. 2). We then integrated the spatially-annotated nonhuman primate data with snRNA-seq data generated from human amygdala tissue to infer the anatomical locations of molecularly-defined human neuron types, and evaluate conservation across species. We found a high degree of conservation of all neuron types across macaques, baboons, and humans (Figs. 3 and 4).

**Figure 1.**
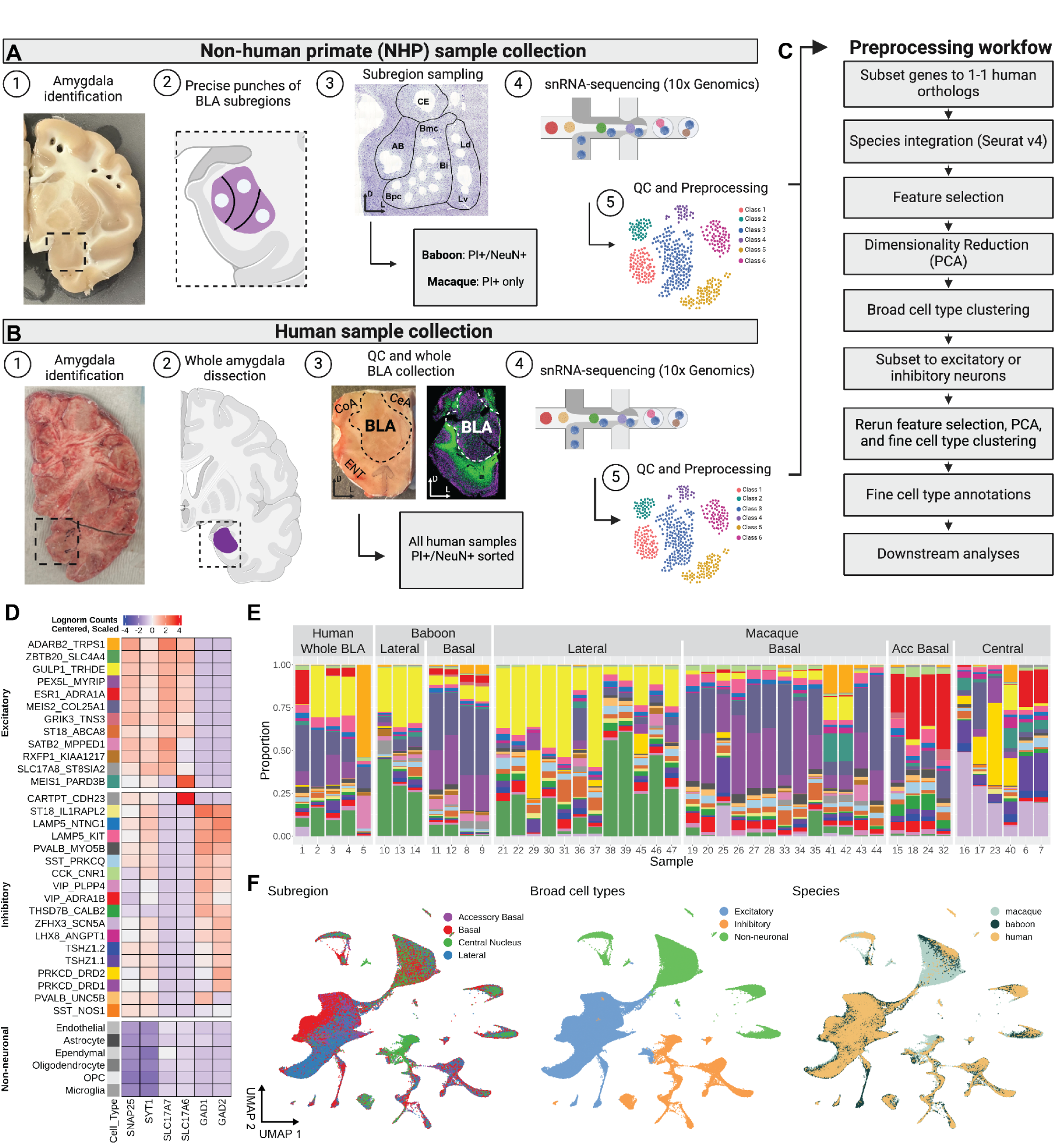
Single-nucleus RNA sequencing (sn-RNAseq) combined with targeted mesoscale sampling to broadly characterize and spatially localize cell types in the amygdala of humans, baboons, and macaques. (A and B) Visualization of targeted mesoscale sampling of one or more amygdala in humans and nonhuman primates (*Macaca mulatta* and *Papio anubis*). The entire basolateral complex in humans including the accessory basal, basal, and lateral nuclei was collected from fresh frozen tissue sections, and all samples were then enriched for neurons using NeuN^+^ staining (81.9% neurons in total). In macaques targeted sampling used biopsy punches of fresh, unfrozen brain slabs to sample the basal, accessory basal, lateral, and central nuclei. In baboons the basal and lateral nuclei were sampled and all samples were then enriched for neurons using NeuN+ staining (80-99% neurons in total). (C) Illustration of the preprocessing workflow to identify cell types across all three species. (D) Heatmap of gene expression for broad marker genes used to characterize cell type clusters as excitatory neurons, inhibitory neurons, or non-neurons. (E) Stacked bar plots depicting the proportion of fine cell types per sample, grouped by species and the amygdala nuclei that were sampled. (F) UMAP visualizations after integration across species of sequenced nuclei colored by amygdala subregion (left), broad cell type annotations (middle), and fine cell type annotations (right). CE, central nucleus; Ld, dorsal region of the lateral nucleus; Lv, ventral region of the lateral nucleus; Bmc, magnocellular region of the basal nucleus, Bi, intermediate region of the basal nucleus; Bpc, parvocellular region of the basal nucleus; AB, accessory basal nucleus.

**Figure 2.**
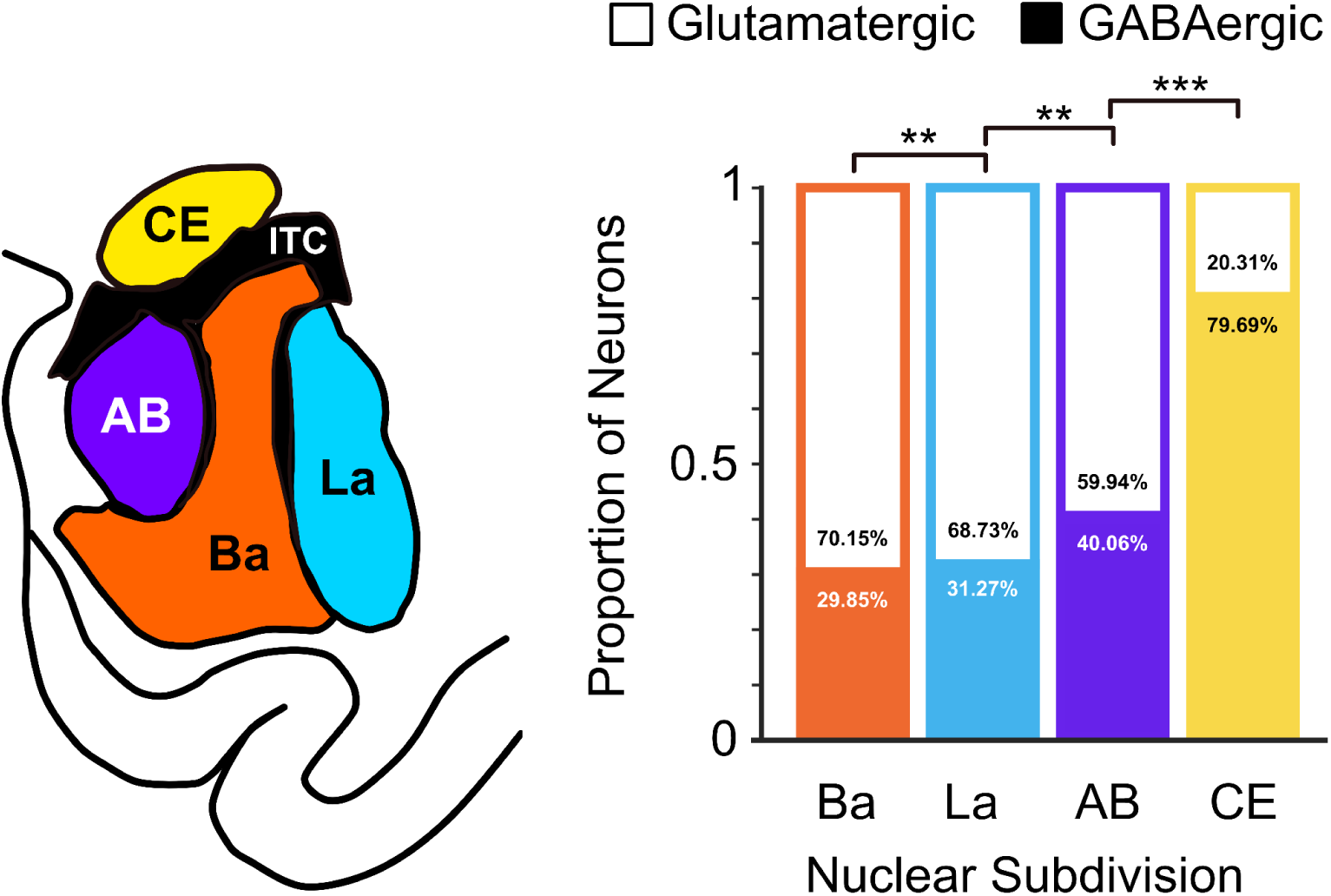
Proportion of neuronal snRNAseq types classified as either glutamatergic or GABAergic from tissue samples of the major amygdala nuclei in macaques. **(A)** Schematic of nuclear subdivisions sampled. **(B)** Each sequenced neuron (*n* = 49,201; Ba = 17,957; La = 16,623; AB = 8,942; Ce = 6,039) was grouped based on its anatomical origins to determine the ratio of glutamatergic to GABAergic neurons in each nuclear subdivision. Pairwise chi-square tests ^35^ demonstrated the proportion of GABAergic neurons differed by subdivision (** *p* < .005; *** *p* < 0.0005). Ba, basal nucleus; La, lateral nucleus, AB, accessory basal; CE, central nucleus.

**Figure 3:**
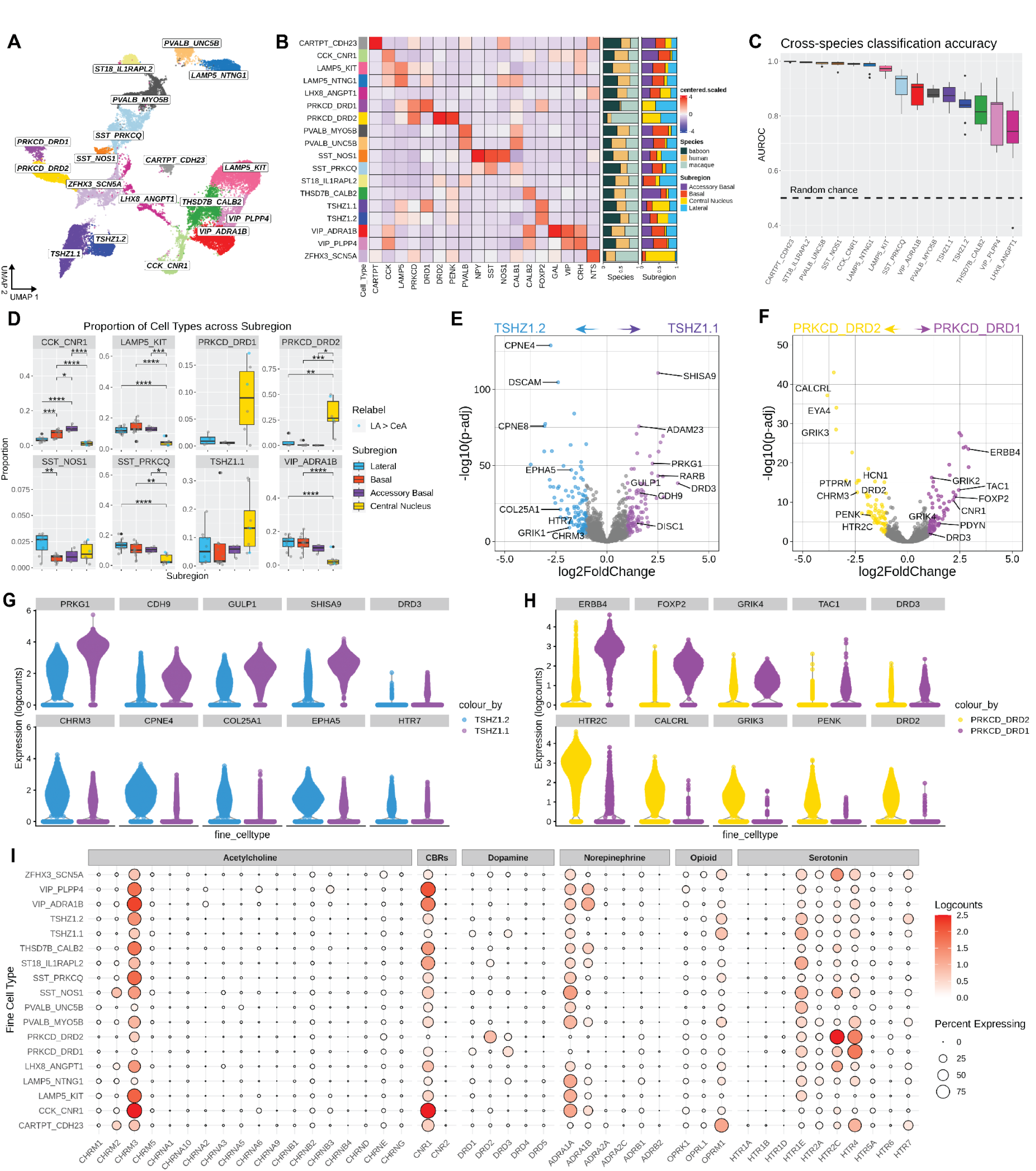
Transcriptomic profiling of inhibitory neurons in the primate amygdala across species and subdivisions. **A**) UMAP visualization colored by inhibitory cell type clusters. **B**) Heatmap showing the mean logcounts of canonical marker gene expression (centered and scaled) across cell type clusters. Stacked bar plots on the right-hand side show the proportion of nuclei in each cell type cluster derived from either nonhuman primate or human species, and the proportion of nuclei derived from each subdivision within macaque samples. **C**) MetaNeighbor area under the receiver operating characteristic (AUROC) values depicting cross-species cell type classification accuracy. AUROC values of 0.5 represent random chance with positive values reflecting above chance accuracy. **D**) Boxplots displaying the proportion of cell types per sample, grouped by subregion punch location in macaque. Two dorsal lateral amygdala (LA) punches were identified as having significant contamination of central amygdala (CeA) cell types (see Methods), and were thus re-labeled as CeA (blue data points) for visualization. Each data point represents one sample. Volcano plots showing pseudobulk differentially expressed genes (DEGs) between the two intercalated cell (ITC) (**E**) and *PRKCD* (**F**) cell type clusters. P-values shown in volcano plots are FDR-adjusted. Colored points represent genes that are significantly upregulated in the respective cell types. Violin plots of the top selected DEGs from the ITC (**G**) and *PRKCD* (**H**) pseudobulk analyses showing the log-transformed counts of nuclei across all three species. **G**) Dot plots showing the mean log-transformed counts (dot color) and percentage of cells expressing (dot size) various genes encoding neurotransmitter and neuromodulator receptors across inhibitory cell types. Box plots display the median with upper and lower quartiles, with whiskers representing the minimum and maximum values. Black data points represent outliers. \**p* < 0.05; \*\**p* < 0.01; \*\*\**p* < 0.001; \*\*\*\**p* < 0.0001 (Tukey’s HSD).

**Figure 4:**
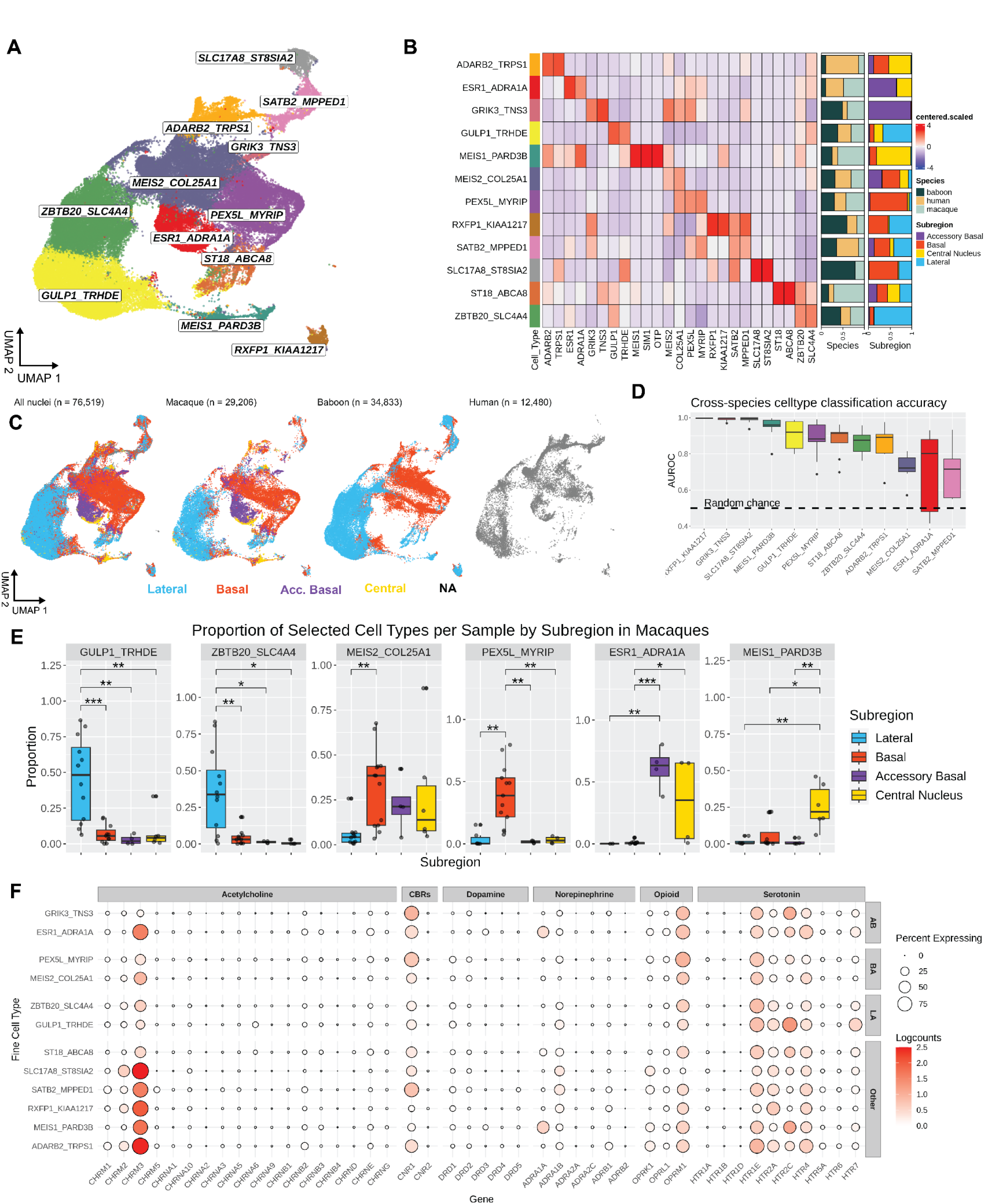
Amygdala excitatory neurons across species and subregions. **A**) UMAP visualization colored by excitatory cell type clusters. **B**) Heatmap showing the mean logcounts of marker gene expression (centered and scaled) across cell type clusters. Stacked bar plots on the right-hand side show the proportion of nuclei in each cell type cluster derived from either non-human primate or human species, and the proportion of nuclei derived from each subregion within macaque samples. **C**) UMAP visualization colored by the amygdala subregion from which nuclei were sampled across species. Nuclei sampled from humans are colored gray (NA) because the entire basolateral amygdala area was sampled rather than precise subregions. **D**) MetaNeighbor AUROC values depicting cross-species cell type classification accuracy. Area under the receiver operating characteristic (AUROC) values of 0.5 represent random chance. **E**) Dot plots showing the mean logcounts (color) and percent of cells expressing (size) various genes encoding neurotransmitter and neuromodulator receptors across inhibitory cell types. Box plots display the median with upper and lower quartiles, whiskers represent min and max. Black data points represent outliers. \**p* < 0.05; \*\**p* < 0.01; \*\*\**p* < 0.001; \*\*\*\**p* < 0.0001 (Tukey’s HSD).

We identified excitatory neuron types that were distinct to the lateral, basal, and accessory basal nuclei and spatially validated neuron type-specific marker gene expression using smFISH analysis in both macaque and human tissue. We also identified two classes of ITC spiny and aspiny interneurons that respectively differed in expression of dopamine receptor D3 (*DRD3*) and serotonin receptor 5HT-7 (*HTR7*). With well-annotated human samples we also examined associations between specific cell types and neuropsychiatric disorders. Multiple classes of GABAergic interneurons, particularly those expressing *LAMP5* and *SST,* were enriched for genes downregulated in postmortem brain tissue of individuals diagnosed with post-traumatic stress disorder (PTSD) ^27^. Moreover, we identified enrichment of genetic risk for several of these disorders across unique neuron types located in the lateral, basal, or accessory basal nucleus.

Our census of molecularly-defined cell types across the five major primate amygdala nuclei is a valuable resource for refining our understanding of structure-function relationships in the amygdala. Incorporating molecular profiling with behavioral and circuit neuroscience approaches has already meaningfully advanced our understanding of amygdala function in rodents ^4–8^. Coupled with existing molecular genetic tools, information in this atlas can enable perturbation studies that selectively target glutamatergic and GABAergic function in specific subdivisions of the primate amygdala to determine their role in motivated behaviors. Moreover, as technological capacities increase, this atlas can be further enriched by addition of molecular profiles for amygdala neurons based on their axonal projection targets, as has been done in mice ^5,28,29^. To facilitate these goals all data and code are freely accessible, and we created cloud-based tools that enable data exploration (see STAR methods).

## RESULTS

### Anatomically informed snRNA-seq data generation from samples of rhesus macaque, baboon and human amygdala

We acquired postmortem tissue containing the amygdala from 5 rhesus macaques (ages 5-18 years, *M* = 11.85; 3 female), 2 female olive baboons (ages 3-4 years, *M* = 3.97; Supplementary Table 1), and 5 human brain donors (ages 34.5-70.64 years, *M* = 50.27; 1 female; Supplementary Table 2). Macaque and baboon brains were dissected into 2-4 mm slabs using a species-specific 3D-printed brain matrix into 1-3 slabs spanning the rostral to caudal extent of the amygdala. Nuclear subdivisions, particularly the basal and lateral nucleus were easily visualized using white matter boundaries. To sample individual nuclei of the amygdaloid complex, we collected targeted tissue punches (1-2.5 mm in diameter) of the accessory basal nucleus, basal nucleus, lateral nucleus, and central nucleus (Fig. 1B). In macaques, tissue punches were taken from all four nuclei, whereas in baboons only the lateral and basal nuclei were sampled, due to the larger more complex folding patterns of the temporal lobe and amygdala. In humans, tissue from the entire basolateral amygdala complex was collected in multiple 100 µm sections from a tissue block containing all major nuclear divisions. Immunostaining to assess gross neuroanatomical structure of the human tissue block and smFISH of canonical marker genes for white matter and excitatory neurons of separate sections were used to visualize and score the boundaries of the basolateral amygdala on the tissue block prior to collecting sections for sequencing (Fig. 1A).

We used the 10x Genomics Chromium platform to generate snRNAseq on the collected nonhuman primate and human samples. Quality control was independently performed for each of the 47 samples (Figs. S1-S3) and preprocessing followed a standard workflow (Fig. 1C) resulting in a final dataset of 171,928 nuclei (>700 UMIs/library; mean = 12,006.78 UMIs). Briefly, we integrated the snRNAseq data first within species (to account for any batch effects) and then across species (based on shared transcriptional signatures across primate genomes), then performed dimensionality reduction to identify broad cell types using canonical marker genes for neuronal (excitatory and inhibitory) and non-neuronal cell types (Fig. 1D). There was a clear dissociation between the central nucleus and the other major nuclei in the overall ratio of GABAergic to glutamatergic neuron types (Fig. 2). Of the neurons that came from tissue punches of the central nucleus 79.6% were GABAergic, whereas neuronal populations derived from punches of the other subdivisions contained at most 40% GABAergic neurons. Glutamatergic neurons were more prevalent in the accessory basal (59.9%), lateral (68.7%), and basal (70.15%) nuclei. This asymmetry in the balance of glutamatergic and GABAergic neurons between amygdalar subdivisions, especially between the central nucleus and basolateral complex, was coarsely assessed across species in prior studies using a variety of methods ^30–34^. However, a direct and unbiased comparison across all cell types and subdivisions was not available until now. This highlights the value of using targeted dissections in combination with snRNAseq to reconstruct the anatomical origins of multiple neuron types.

To delve further into how specific subtypes of GABAergic and glutamatergic neurons were distributed across the four major nuclei, we separately subset glutamatergic and GABAergic neurons from other cell types into two independent datasets. We then repeated feature selection, dimensionality reduction, clustering and cell type annotation to derive fine cell types. Overclustering was avoided using the pairwise modularity ratio of observed to expected edge weights to ensure all clusters were well separated. This analysis identified 18 types of GABAergic neurons and 12 types of glutamatergic neurons (Fig. 1D). When we examined the frequency of each cell type within each sample organized by species and location we observed that all cell types were present in all three species (Fig. 1E; left panel, 1F). However, specific cell types and amygdala nuclei appeared to be linked when each datapoint in our dimensionality reduction plots was relabeled according to locations sampled in nonhuman primates (middle panel, Fig. 1F). Even at this coarse level, it was apparent that multiple glutamatergic neuron types were specific to either the lateral or basal nucleus, as well as multiple GABAergic neuron types specific to the central nucleus (right panel, Fig. 1F).

### GABAergic neuron types in the primate amygdala show distributed or subregion specific expression patterns

The broad types of GABAergic neurons found in the primate amygdala are well conserved and extensively characterized ^15,36^. Our intent was to molecularly profile each type, determine the degree of heterogeneity (i.e. number of subtypes), and whether neuron types were uniquely expressed in a nuclear subdivision. With the expectation that our targeted dissections should include neurons from the intercalated cell masses that fill the spaces in between the other major nuclei, we were particularly interested in using *TSHZ1* and *FOXP2* as primate-conserved markers of ITC neurons to derive the number of types that existed and their complete molecular profiles. We were also keen to molecularly profile protein kinase C-δ^+^(*PKRCD^+^*) interneurons in the central nucleus considering their important role in mediating fear and anxiety in rodent models ^1,37^, and the relative lack of information about their function in the primate amygdala ^38^.

Neurons were labeled as GABAergic if they co-expressed *SNAP25*, *SYT1*, or *RBFOX3* in combination with *GAD1* or *GAD2*. To enable a more detailed analysis of the different inhibitory neuron classes we first subset all of the inhibitory neurons from the rest of the dataset and repeated feature selection, dimensionality reduction, and clustering. We identified 18 different types of inhibitory neurons in the primate amygdala (Fig. 3A) with representation of all major interneuron classes (*SST*, *PVALB*, *VIP*, *CCK*, and *LAMP5*). We also identified clusters corresponding to protein kinase C-δ^+^(*PKRCD^+^/SST^-^*) interneurons in the central nucleus, as well as interneurons expressing the cocaine and amphetamine regulated transcript (CART) peptide ^39^ and neurotensin. Inhibitory neuron clusters were broadly organized by expression of transcription factors corresponding to their developmental origins in the lateral (LGE; *MEIS2^+^*, *ZFXH3^+^*), medial (MGE; *ADARB2^+^*, *PROX1^+^*, *NPAS3^+^*, *NR2F1^+^*), or caudal (CGE; *LHX6^+^*, *SOX6^+^*) ganglionic eminence. *SST^+^*, *PVALB^+^*, and *LAMP5^+^* interneurons expressed MGE transcriptomic factors, whereas *VIP^+^* and *CCK^+^* interneurons expressed CGE transcriptomic factors. *PKRCD^+^* and *CART^+^* inhibitory neurons expressed LGE transcription factors.

#### VIP and CCK interneurons

*VIP^+^* interneurons tend to be disinhibitory cells that largely target other interneurons types such as *SST^+^* and *PVALB^+^* interneurons, and in the rodent amygdala are critical for associative learning ^40^. We found two clusters of VIP^+^ interneurons (*VIP^+^/ADRA1B^+^*and *VIP^+^/PLPP4^+^*), which represented 9.71% and 3.47%, respectively, of all inhibitory neurons analyzed. Compared to other GABAergic populations, both clusters showed increased expression of genes (Fig. 3B) encoding calretinin (*CALB2*), cholecystokinin (*CCK*), corticotropin releasing hormone (*CRH*), cannabinoid receptor 1 (*CNR1*), the M3 acetylcholine receptor (*CHRM3*; Fig. 3I), and multiple alpha-1-adrenoceptors (*ADRA1A* and *ADRA1B*; Fig. 3I) that bind norepinephrine. The *VIP^+^/ADRA1B^+^*cluster was further characterized by enriched expression of the gene encoding the neuropeptide galanin (*GAL*; Fig. 3B). While galanin immunoreactivity is detected in the nonhuman primate amygdala ^41^, it is not directly linked to *VIP* interneuron function. For each cluster we also found that in macaques, a greater proportion of the *VIP^+^/CCK^+^* neurons came from tissue punches of either the accessory basal, basal, or lateral nuclei than tissue punches of the central nucleus (Fig. 3D and Fig. S6), despite the central nucleus containing an overall greater proportion of GABAergic neurons (Fig. 2).

A third cluster of *CCK^+^* neurons (*CCK^+^/CNR1^+^*) consisted of 5.28% of all inhibitory neurons analyzed. The neurons in this cluster had a similar gene expression profile to the other two *VIP^+^*clusters (Fig. 3D), but were distinguished by expression of the cannabinoid receptor 1 gene (*CNR1;* Fig. 3I). A greater proportion of the neurons in this cluster were also present in samples of the accessory basal or basal nucleus compared to either the lateral or central nucleus in macaques (Fig. 3D). This *CCK^+^/CNR1^+^* cluster is likely homologous to mouse *CCK^+^* basket cells, which synapse onto the soma and proximal dendrites of excitatory neurons in the ^42^ amygdala ^15,36^.

#### SST interneurons

Somatostatin-containing (*SST+*) interneurons are dendritic-targeting cells that regulate afferent inputs and synaptic plasticity, and recent work in rodents suggests that amygdalar *SST*^+^ interneurons play a critical role in gating the retrieval of associative memories ^42^.We identified a large cluster of *SST^+^* interneurons (*SST^+^/PRKCQ^+^*) representing 9.5% of all inhibitory neurons analyzed. Neurons in this cluster showed relatively increased expression of calbindin (*CALB1*) and decreased expression of nitric oxide synthase 1 (*NOS1)*. This aligns with the typical gene expression profile of *SST*-containing interneurons in the rodent amygdala ^15,36,43^. In macaques, neurons in this cluster were on average more prevalent in samples of either the accessory basal (10.35%), basal (10.03%), or lateral nucleus (13.73%) in comparison to samples of the central nucleus (3.92%; Fig. 3D).

A second and smaller *SST^+^* cluster of inhibitory neurons (*SST^+^/NOS1^+^*) consisted of 1.4% of all inhibitory neurons analyzed. The neurons in this cluster showed enriched expression of neuropeptide Y (*NPY^+^*), *NOS1^+^*, and chondrolectin (*CHODL^+^*), but depletion of calbindin (*CALB1*) expression. Co-expression of *SST* and *NOS1* suggests that these neurons are GABAergic projection neurons, which have been previously described in the rodent amygdala, hippocampus, and entorhinal cortex ^32,36,44^. In addition, this cluster showed increased expression of genes encoding the M2 and M3 acetylcholine receptors (*CHRM2* and *CHRM3*), alpha-1-adrenoceptor (*ADR1A*), and multiple serotonin receptors (*HTR1E, HR2C, and HTR4*). GABAergic projection neurons in the rodent amygdala that are similarly immunoreactive for SST, NPY, and the M2 acetylcholine receptor project to the entorhinal cortex ^45,46^ and immunohistochemical characterizations indicate that they are spatially-localized near white matter tracts running through the basal and lateral nuclei ^32,36^. Accordingly, we found a greater proportion of *SST^+^/NOS1^+^* neurons in tissue punches from the lateral (2.28%) compared to the basal nucleus (0.9%; Fig. 3D).

#### PVALB interneurons

Amygdalar *PVALB*^+^ interneurons are known to coordinate oscillations to drive both behavioral state transitions ^47–49^ and play critical roles in the encoding and retrieval of associative memories ^50,51^. Prior immunohistochemical studies in the macaque amygdala suggest the presence of two subtypes of PVALB-containing interneurons: basket cells and chandelier cells ^36^. We identified two *PVALB^+^*inhibitory neuron clusters, *PVALB^+^*/*UNCB5^+^* and *PVALB^+^*/*MYO5B^+^*, that respectively represented of 4.46% and 6.17% of all inhibitory neurons analyzed. Relative to other interneurons, both clusters showed increased expression of *CALB1^+^*. *UNCB5* encodes a receptor for netrins, which are secreted molecules that regulate axonal growth and angiogenesis, and *MYO5B* encodes myosin Vb, which transports recycling endosomes in dendritic spines to support endocytosis of AMPA receptors and synaptic plasticity. Marker gene expression suggests that the *PVALB^+^/UNCB5^+^* cluster are chandelier interneurons and *PVALB^+^/MYO5B^+^* cluster are basket cells, but these assignments need to be verified with additional morphological data. However, doing so should be of interest because these two clusters differed in their spatial localization. In macaques, *PVALB^+^/UNCB5^+^* interneurons were more prevalent in samples of the accessory basal, basal, or lateral nucleus compared to the central nucleus, while a larger proportion of *PVALB^+^*/*MYO5B^+^* were found in samples taken from the basal nucleus compared to either the lateral or central nucleus (Fig. S6).

We identified two additional clusters (*ST18^+^*/*IL1RAPL2^+^*and *CARTPT^+^*/*CDH23^+^*) that relative to other inhibitory neurons showed increased *PVALB* expression, and no overlapping expression with other canonical interneuron markers. The *ST18^+^*/*IL1RAPL2^+^* cluster consisted of 2.8% of the inhibitory neurons analyzed. The *ST18^+^*/*IL1RAPL2^+^* cluster is noteworthy because *IL1RAPL2* and *IL1RAPL1* are located at a region on chromosome X associated with autism ^52^. The function of *IL1RAPL2* is unexplored, but *IL1RAPL1* contributes to synapse stabilization and maintenance ^53^. The *ST18^+^*/*IL1RAPL2^+^* cluster also exhibited enriched expression of the neurokinin B receptor (*TACR3^+^*) and neurokinin B receptor agonism in the basolateral amygdala increases fear potentiated startle ^54^. Amongst inhibitory neurons, *IL1RAPL2^+^* neurons also showed the highest expression of *HTR1E* relative to all other inhibitory neurons, suggesting sufficiency for serotonergic modulation. The majority of the neurons in the *ST18^+^*/*IL1RAPL2^+^* cluster were derived from tissue punches taken from the lateral nucleus (Fig. S6).

Co-expression of *PVALB* and the neuropeptide *CART* (cocaine and amphetamine regulated transcript) in interneurons is well-established ^55^. The *CARTPT^+^*/*CDH23^+^* cluster consisted of 1.2% of all the inhibitory neurons analyzed. Consistent with known expression patterns of CART in the primate amygdala equivalent proportions of neurons were sampled from all four of the major nuclei (Fig. S6).

#### LAMP5 interneurons

Neurogliaform cells in the amygdala are medium-sized GABAergic neurons with dense perisomatic axonal arborization ^15^. Most axon terminals of neurogliaform cells do not form synapses. Instead they oppose the dendrites of excitatory neurons and release GABA into the extracellular space. In the cortex, neurogliaform cells expressing *LAMP5* are enriched in primates as compared to mice ^56^. We identified two clusters of *LAMP5^+^* interneurons, *LAMP^+^*/*KIT^+^*and *LAMP5^+^*/*NTNG1^+^*. Respectively, the clusters represented 10.86% and 5.97% of all the inhibitory neurons analyzed. Both clusters were highly conserved across the three primate species and co-expressed *CCK^+^*. In both clusters, a greater proportion of neurons came from samples of the accessory basal, basal, or lateral nucleus relative to the central nucleus (Fig. 3B). The clusters differed in expression of muscarinic acetylcholine and serotonin receptors (Fig. 3I). The *LAMP^+^*/*KIT^+^*cluster showed enriched expression of the gene encoding the M3 acetylcholine receptor (*CHRM3*) and minimal expression of serotonin receptor genes. By comparison the *LAMP5^+^*/*NTNG1^+^* cluster showed minimal expression of *CHRM3* and relatively increased expression of multiple serotonin receptor genes (*HTR1E*, *HTR2A*, *HTR2C*, *HTR4*).

#### Neurotensin-containing interneurons

Neurons that release the peptide neurotensin (*NTS*) are prevalent in the central nucleus and ITCs in both rodents and primates. We identified two inhibitory neuron clusters with relatively high expression of *NTS*. The *ZFHX3^+^/SNC5A^+^* cluster, representing 7.9% of all the inhibitory neurons analyzed, showed the highest expression of *NTS* across all of the inhibitory neuron clusters. Otherwise these neurons showed similar, albeit weaker, expression of the same marker genes that were enriched in the *PKRCD^+^*clusters (i.e. *PRKCD*, *PENK*, and *CALB1*). Hence, it was unsurprising that this cluster originated almost exclusively from tissue punches of the central nucleus of macaques (Fig. S6). Since we did not systematically sample the central nucleus in baboons and humans, we did not have enough *NTS^+^* neurons to quantitatively assess whether this cell type was conserved across species, but we did however detect it in both species (Fig. 3A and C).

The *LHX8^+^/ANGPT1^+^* cluster consisted of 3.74% of all inhibitory neurons analyzed and did not show increased expression of any canonical interneuron markers besides *GAD1* and *GAD2* (Fig. 1D). However, these neurons did show increased expression of *NTS*, and were conserved across all three species, albeit at a lower level than other cell types. The cluster contained equivalent proportions of neurons sampled from all four major nuclei.

### Intercalated cell masses contain spiny and aspiny interneurons that differ in their expression of dopamine, serotonin, and cholinergic receptors

The ITC masses are relatively understudied in primates relative to rodents, despite their critical role in amygdala function ^1,25,57–60^. We found two inhibitory neuron clusters that co-expressed *TSHZ1* and *FOXP2*, which are established molecular markers of ITCs in rodents (*TSHZ1.1*^+^/*FOXP2^+^* and *TSHZ1.2^+^*/*FOXP2^+^*). Both clusters were also characterized by expression of the transcription factors *MEIS2* and *PBX3,* and the genes encoding the D1 dopamine receptor (*DRD1*) and μ-opioid receptor (*OPMR1*), especially the larger *TSHZ1.1^+^*/*FOXP2^+^* cluster (Fig. 3I). These findings align with prior immunohistochemical and molecular characterizations of ITC neurons ^25,26^.

ITC neurons are localized to either the medial paracapsular cluster, lateral paracapsular cluster, or main islands. These clusters are diffuse, especially in primates ^26^, and they form an extensive neuronal net that fills in the space between the borders of all the major amygdalar nuclei. Neuronal density, however, is highest in the space between the dorsal boundaries of the basal, lateral, and central nucleus. To determine the spatial distribution of the *TSHZ1^+^* clusters we compared the proportion of *TSHZ1^+^* inhibitory neurons in each cluster derived from tissue punches of the four nuclei sampled in macaques. As expected, the proportion of interneurons grouped into the two *TSHZ1^+^*clusters did not differ between the four nuclei sampled (Fig. S6). Although on average and within the larger *TSHZ1.1^+^/FOXP2^+^* cluster, tissue punches of the central nucleus contained a greater proportion of *TSHZ1^+^* inhibitory neurons than lateral, basal, or accessory basal nuclei (Fig. 3D).

To determine how the two *TSHZ1^+^*/*FOXP2^+^*clusters differed from one another we conducted a pseudobulk analysis to identify genes that were differentially expressed in each cluster (Fig. 3E and G). A total of 104 genes were enriched in the larger *TSHZ1.1^+^/FOXP2^+^* cluster (8.16% of all inhibitory neurons analyzed). This included *DRD3* and a gene (*DISC1*) that has been implicated in several neuropsychiatric disorders ^61^. Additional upregulated genes also influenced signaling pathways (*PDE7B, PRKG1, GULP1*), cell adhesion (*CDH9, CDH11, CDH12, FAT1, NECTIN*), cytoskeletal organization (*ACTN2, ARHGAP6, ARHGAP10*), and neuronal development (*BCL11B*, *RARB*), as indicated by genome-scale phylogenetics ^62^.

A total of 115 genes were enriched in the smaller *TSHZ1.2^+^/FOXP2^+^* cluster (4.23% of all inhibitory neurons analyzed). These included genes encoding serotonin (*HTR7*), muscarinic cholinergic (*CHRM3*), glutamate (*GRIK1*, *GRIA4*), and GABA receptors (*GABRG1*), and a glutamate receptor interacting protein involved in anchoring glutamate receptors at synaptic sites (*GRIP2*; Fig. 3E, G). Interestingly, this cluster also showed enrichment for the transcription factor *PAX6* which is a marker of the main islands of the ITC ^25^. Additional upregulated genes influenced signal transduction pathways (*TGFB2, ALK, EPHA5*), neuronal excitability (*KCNJ16, KCNQ1*), axon guidance (*NTNG1,SLIT2, DCC, CNTN3, SEMA3E, SEMA6D*), extracellular matrix organization (*COL25A1, FBN2, LAMA4, and LTBP1*), cell adhesion (*CDH13, PCDH15*), and neural development (*RELN, NPAS3*), as indicated by genome-scale phylogenetics ^62^.

A prior examination of ITC neurons in nonhuman primates identified at least two morphological types of GABAergic neurons, small spiny neurons and large aspiny neurons ^26^. Our transcriptomic profiling of ITCs affirms this classification. Enriched expression of the transcription factor *BCL11B* and the retinoic acid receptor *RARB* in the larger ITC cluster (*TSHZ1.1^+^/FOXP2^+^*) is of interest because these genes regulate development of medium spiny neurons in the striatum ^63^. This implies that the larger *TSHZ1.1^+^/FOXP2^+^* cluster consists of dopaminoceptive spiny neurons expressing D1 and D3 receptors. Whereas enriched expression of *RELN* in the smaller ITC cluster (*TSHZ1.2^+^/FOXP2^+^*) suggests it contains aspiny neurons that are receptive to multiple neurotransmitters through expression of D1 receptors, 5HT-7 serotonin receptors, and M3 acetylcholine receptors.

### Protein kinase C-δ interneurons in the nonhuman primate central nucleus differ based on dopamine receptor expression

Protein kinase C-δ (*PKRCD*) interneurons are located in the central nucleus where they play a critical role in amygdala-dependent associative learning through parallel interactions with SST^+^ interneurons, nociceptive inputs from the parabrachial nucleus, and projections to the dopaminergic midbrain ^1,36,37^. We identified two clusters of *PKRCD* interneurons, which were distinguished by differential expression of dopamine and kainate receptors (*PRKCD^+^*/*DRD1^+^* and *PRKCD^+^*/*DRD2^+^*; Fig. 3F). Both clusters expressed high levels of *HTR4* and *HTR2C,* especially relative to other inhibitory cell types (Fig. 3I). The larger *PRKCD^+^*/*DRD2^+^*cluster represented 8.3% of all inhibitory neurons analyzed while the smaller *PRKCD^+^*/*DRD2^+^* cluster contained 2.52%. Both clusters were almost exclusively derived from central and lateral nuclei tissue samples (Fig. 3D). Since it is well-established that *PKRCD* neurons are located in the central nucleus, the neurons in each cluster that originated from the lateral nucleus were likely from dorsal punches of the lateral nucleus which unintentionally included bordering tissue from the central nucleus.

Prior evaluations of *PKRCD^+^* interneuron function identified the importance of D2 receptor expression ^37,64^. Despite reporting that neurons in the central nucleus expressed either D1 or D2 receptors, previous studies did not examine overlap between *PKRCD* and *DRD1* expression ^37^. To further investigate how the two *PKRCD^+^* interneuron clusters differed from one another we conducted a pseudobulk analysis to identify genes that were differentially expressed in each cluster. In addition to increased expression of *DRD2*, the *PRKCD^+^*/*DRD2^+^* cluster showed enriched expression of genes encoding the 5HT-2C serotonin (*HTR2C*), M3 cholinergic (*CHRM3*), calcitonin (*CALCR*), netrin (*UNC5D*), and glutamate kainate (*GRIK3*) receptors. Expression was also enriched for the endogenous opioid peptide proenkephalin (*PENK*). This gene expression profile aligns with functional characterizations of *PKRCD^+^* interneurons in rodents ^37,65^.

Transcriptomic profiling of the *PRKCD^+^*/*DRD1^+^* cluster, however, indicated it might represent a subtype of *PRKCD+* interneurons that are related to ITC neurons. Relative to its counterpart, the *PRKCD^+^/DRD1^+^*cluster showed enriched expression of the ITC marker gene *FOXP2*, as well as increased expression of the gene encoding the D3 receptor (*DRD3*). Therefore this cluster is more similar to the *TSHZ1.1+/FOXP2+* cluster that also showed enriched D1 and D3 receptor expression and was more prevalent in the central nucleus. Additional upregulated genes in the *PRKCD^+^/DRD1^+^* cluster included glutamate kainate receptors (*GRIK2* and *GRIK4*), transmembrane receptors involved in axon guidance and synapse formation (*ERBB4* and *ROBO2*), and the endogenous opioid peptide prodynorphin (*PDYN*; Fig. 3F).

### Primate amygdala contains multiple types of excitatory neurons that are organized by subregion

Compared to GABAergic neurons, clear hypotheses about the number of glutamatergic neuron types in the primate amygdala are lacking. Prior neuroanatomical studies noted that glutamatergic neurons differ in terms of their size (e.g. magnocellular versus parvocellular subdivisions particularly in the basal nucleus) and excitability ^15,43,66^, but have not identified neuron types specific to nuclear subdivisions. We adopted the same approach that we took in detailing types of GABAergic neurons. After subsetting all of the neurons labeled as glutamatergic that co-expressed *SNAP25*, *SYT1*, or *RBFOX3* in combination with *SLC17A7* (VGLUT1) or *SLC17A6* (VGLUT2), we repeated feature selection, dimensionality reduction, and clustering using the gene expression profiles of just glutamatergic neurons. We identified 12 types of glutamatergic neurons in the primate amygdala (Fig. 4A and B). Expression of either *SLC17A7* or *SLC17A6* (Fig. 1D) suggests that these clusters represent pyramidal-like projection neurons ^15^.

When examining the UMAP visualizations for each species with each neuron labeled by amygdala subregion of origin, we observed anatomical segregation of excitatory neuron cell types with several cell types mapping to only a single nucleus. For clusters mapping to more than one nucleus, distribution across the four major nuclei aligned with established expression patterns of particular receptors for major neurotransmitters or hormones ^15^. Moreover, there was complete overlap between human excitatory neurons whose anatomical origins were unknown and the nonhuman primate excitatory neurons whose anatomical origins were well-defined (Fig. 4C). A decoding analysis across species confirmed that the neuron types were highly conserved across nonhuman primates and humans (Fig. 4D). Therefore, for clusters of excitatory neurons with a clear anatomical origin in nonhuman primates, we annotated each of the human excitatory neurons in those clusters with the same label. This finding illustrates the utility of using targeted mesoscale tissue dissections in nonhuman primates to recover the spatial identity of cells in regions of the human brain where precise anatomical dissections are not possible.

We identified four clusters of excitatory neurons (*RXFP1^+^/KIAA127^+^, ADARB2^+^/TRPS1^+^, SLC17A8^+^/ST8SIA2^+^,* and *SATB2^+^/MPPED1^+^*) that were not overrepresented within a particular nuclear subdivision. Instead, neurons in these clusters originated equally from two or more nuclear subdivisions (Fig. S11) and altogether they represented 23.1% of the excitatory neurons analyzed. These neuron types showed increased expression of muscarinic and nicotinic acetylcholine receptors (Fig. 4F), particularly genes encoding the M2 or M3 acetylcholine receptor (*CHRM2* and *CHRM3*). In macaques, the neurons in these clusters were predominantly derived from tissue punches from the basal nucleus, but also included neurons from the accessory basal nucleus, lateral nucleus, and central nucleus (Fig. S6). In primates, the magnocellular division of the basal nucleus, followed by all other subdivisions of the basal nucleus, the magnocellular and superficial subdivisions of the accessory basal nucleus, the ventrolateral region of the lateral nucleus, and the central nucleus all receive substantial cholinergic input from the basal forebrain. Interestingly, all of these cell types showed stronger expression of the M3 cholinergic receptor gene *CHRM3* compared to the type 1 muscarinic cholinergic receptor (*CHRM1*), which has been the focus of most prior studies in rodents and primates ^45,67^.

### Glutamatergic neuron types specific to nuclear subdivisions

We identified 7 additional clusters of excitatory neurons that were localized to a particular nucleus of the amygdala (Fig. 4B and C). We identified two clusters of excitatory neurons that came almost exclusively from the accessory basal nucleus: *ESR1^+^/ADRA1A^+^* and *GRIK3^+^/TNS3^+^.* The *ESR1^+^/ADRA1A^+^* cluster consisted of 21.1% of the excitatory neurons analyzed, whereas the *GRIK3^+^/TNS3^+^* cluster contained 1.1% of the excitatory neurons. Cross-species decoding analyses indicated that the *GRIK3^+^/TNS3^+^* neuron type was more conserved than *ESR1^+^/ADRA1A^+^*neuron type. In any case, our results are consistent with prior neuroanatomical findings in macaques and humans that estrogen receptor expression is highest in the accessory basal nucleus ^68–70^.

Our approach was also successful in identifying glutamatergic neuron types specific to either the basal or lateral nucleus (Fig. 4A and C). We identified two clusters of excitatory neurons that came almost exclusively from tissue punches of the lateral nucleus in macaques and baboons: *GULP1^+^*/*TRHDE^+^* and *ZBTB20^+^*/*SLC4A4*. These clusters, respectively, contained 14.26% and 9.59% of all the excitatory neurons analyzed. We detected increased expression of the top marker genes *GULP1*, *ZBTB20*, and *SLC4A4* in both clusters, suggesting quantitative rather than qualitative differences in the transcriptomic profiles of each cell type. However, we did find increased expression of the genes encoding the 5-HT2C and 5-HT7 serotonin receptors in the *GULP1+* cluster. Neurons in both clusters were much more prevalent in samples of the lateral nucleus compared to other amygdala nuclei (Fig. 4E).

We also identified a cluster of excitatory neurons that came almost exclusively from tissue punches of the basal nucleus in macaques and baboons: *PEX5L^+^*/*MYRIP^+^*. This cluster contained 10.67% of all excitatory neurons analyzed and was notable for enriched expression of *CNR1* and *HTR1E* (Fig. 4F). A second, larger cluster, *MEIS2^+^*/*COL25A1^+^*,was populated by neurons that came mostly from either the accessory basal, basal, or central nuclei, but was notable because it contained very few neurons from the lateral nucleus (Fig. 4E). The *MEIS2^+^*/*COL25A1^+^*cluster contained 19.53% of the glutamatergic neurons analyzed. Overall the two clusters exhibited similar neuromodulatory gene expression profiles, and we detected increased expression of *COL25A1* in both neuron types (Fig. 4B).

Having identified marker genes for multiple neuron types that were specifically located in either the basal or lateral nucleus, we reasoned that aggregating these neuron types based on their anatomical origins would allow us to identify genetic markers of glutamatergic neurons selectively expressed in these two major subdivisions of the primate amygdala. Because of known issues in interpreting differential gene expression based on UMAP representations we used pseudobulk analyses with the a priori labels assigned to neurons derived from samples of either the accessory basal, basal, or lateral nucleus in macaques and baboons and the post hoc anatomical labels assigned to neurons in the corresponding clusters in samples of the basolateral complex of humans.

Principal component analysis of nonhuman primate pseudobulked samples confirmed that both species and amygdalar subdivision were top sources of variability in gene expression (Fig. S12A). Pseudobulk differential expression analysis detected 147 genes that were upregulated in glutamatergic neurons localized to the lateral nucleus and 219 genes that were upregulated in the basal nucleus. In these differentially expressed gene sets, *GULP1* and *COL25A1* were among the top marker genes for excitatory neurons in the lateral and basal nuclei, respectively, of nonhuman primates (Fig. 5A and B). We next investigated whether these marker genes would be conserved in the human amygdala. To test this, we first collapsed the fine cell type clusters derived largely of lateral origin (*GULP1+*/*TRHDE+* and *ZBTB20+*/*SLC4A4*) and basal origins (*MEIS2+*/*COL25A1+* and *PEX5L+*/*MYRIP+*) into putative superclusters. Within species, we then performed differential expression analysis to get species-specific marker genes for putative lateral, basal, and accessory basal (*ESR1+*/*ADRA1A+*) neuron types and correlated the log-fold change between human and macaques to discover conserved markers of neurons derived from either the lateral (Fig. 5C), basal (Fig. 5D), or accessory basal nucleus (Fig. 5E). This again identified *GULP1*, *COL25A1, ESR1* as the top conserved markers for predicting localization of an excitatory neuron to particular amygdala subdivisions.

**Figure 5:**
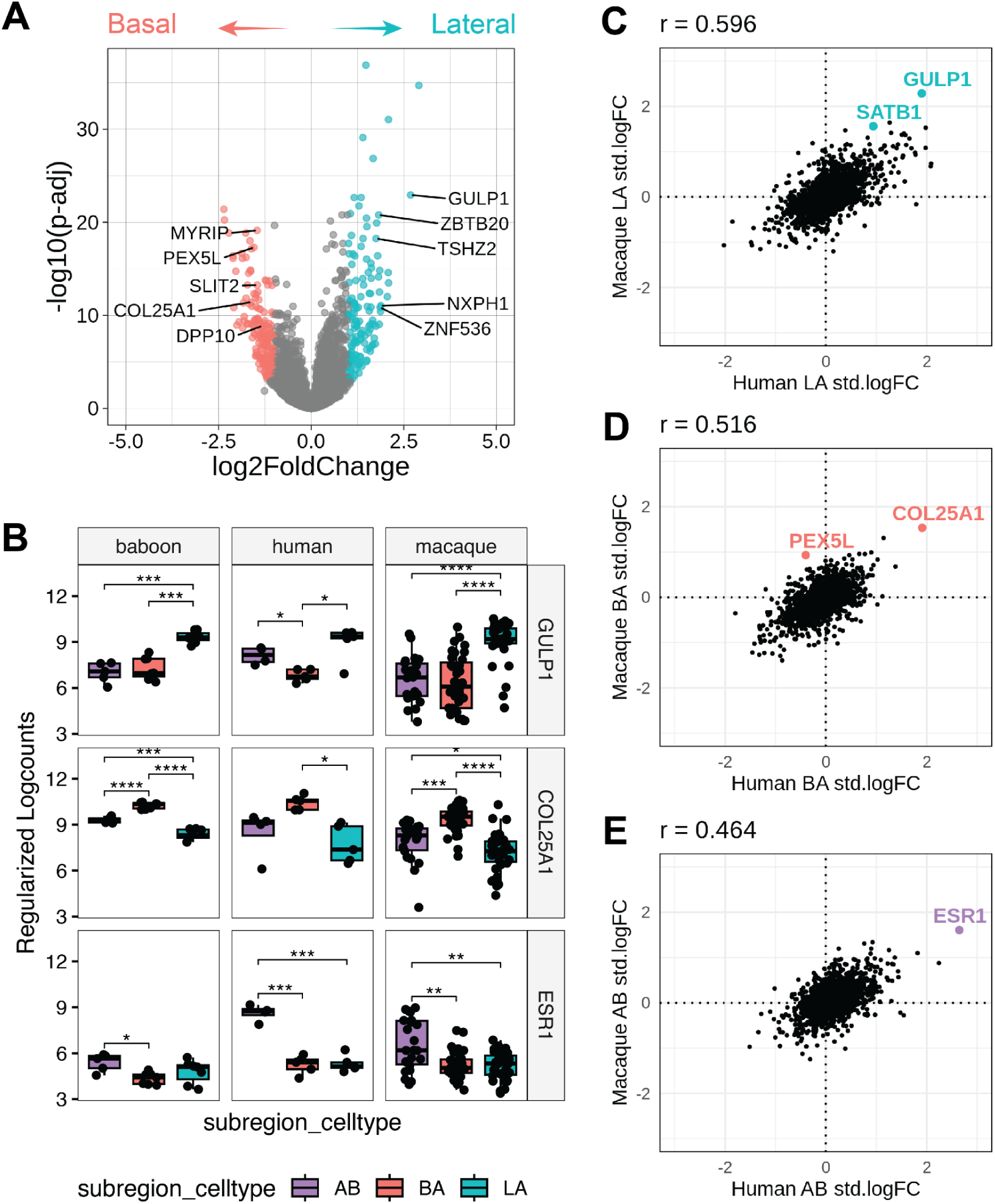
Pseudobulk differential expression analysis of lateral versus basal excitatory neurons reveals conserved marker genes and molecular function. (**A)** Volcano plot showing differentially expressed genes (DEGs) between pseudobulk excitatory neurons derived from either the basal amygdala (BA, salmon) or lateral amygdala (LA, turquoise) subregions in non-human primates. The top five candidate marker genes are shown for each subregion after filtering out lowly expressed genes. P-values shown are FDR-adjusted. **(B)** Box plots comparing mean regularized expression of putative marker genes across subregion clusters for each species. Data points represent individual pseudobulk samples. Scatter plots of human vs macaque standard log2 fold change for putative subregion clusters demonstrate that *GULP1*, *COL25A1*, and *ESR1* are the top conserved marker genes for LA **(C)**, BA **(D)**, and AB **(E)** subregions, respectively. Box plots display the median with upper and lower quartiles. AB, accessory basal, BA basal, LA lateral. \**p* < 0.05; \*\**p* < 0.01; \*\*\**p* < 0.001; \*\*\*\**p* < 0.0001.

To further establish these genes as high-confidence markers of spatially distinct glutamatergic cell types across primate species, we assessed spatial patterns of *GULP1* and *COL25A1* expression among excitatory neurons co-expressing *SLC17A7* using smFISH in tissue from independent macaques (*n* = 2; 2 female) and a human donor (*n* = 1; 1 male) (Fig. S15). As predicted from our cross-species transcriptomic profiling using tissue punch annotations, there was no overlap in the spatial expression of these marker genes in the macaque (Fig. 6 and Fig. S16). Using myelin basic protein (*MBP*) gene expression to define boundaries between the major amygdala nuclei, we found *GULP1* expression was confined to the lateral nucleus, whereas *COL25A1* expression was present throughout all subdivisions of the basal nucleus and also present in the accessory basal nucleus (Fig. 5A-C and Fig. S16). In humans, we similarly found that *COL25A1* expression was restricted to the basal nucleus and that *GULP1* expression predominated in the lateral nucleus (Fig. 6D-F). Compared to macaques, we did observe more spatial overlap in the gross expression of *GULP1* and *COL25A1* in parts of the basal nucleus in humans, but we did not observe co-expression of these two genes within individual *SLC17A7^+^*nuclei. This is somewhat expected (i.e. neither cluster of sequenced, excitatory neurons was absolutely derived from one nuclear subdivision) and the boundaries between the lateral and basal nuclei in humans are more difficult to define given the complex and irregular architecture of the human amygdala.

**Figure 6.**
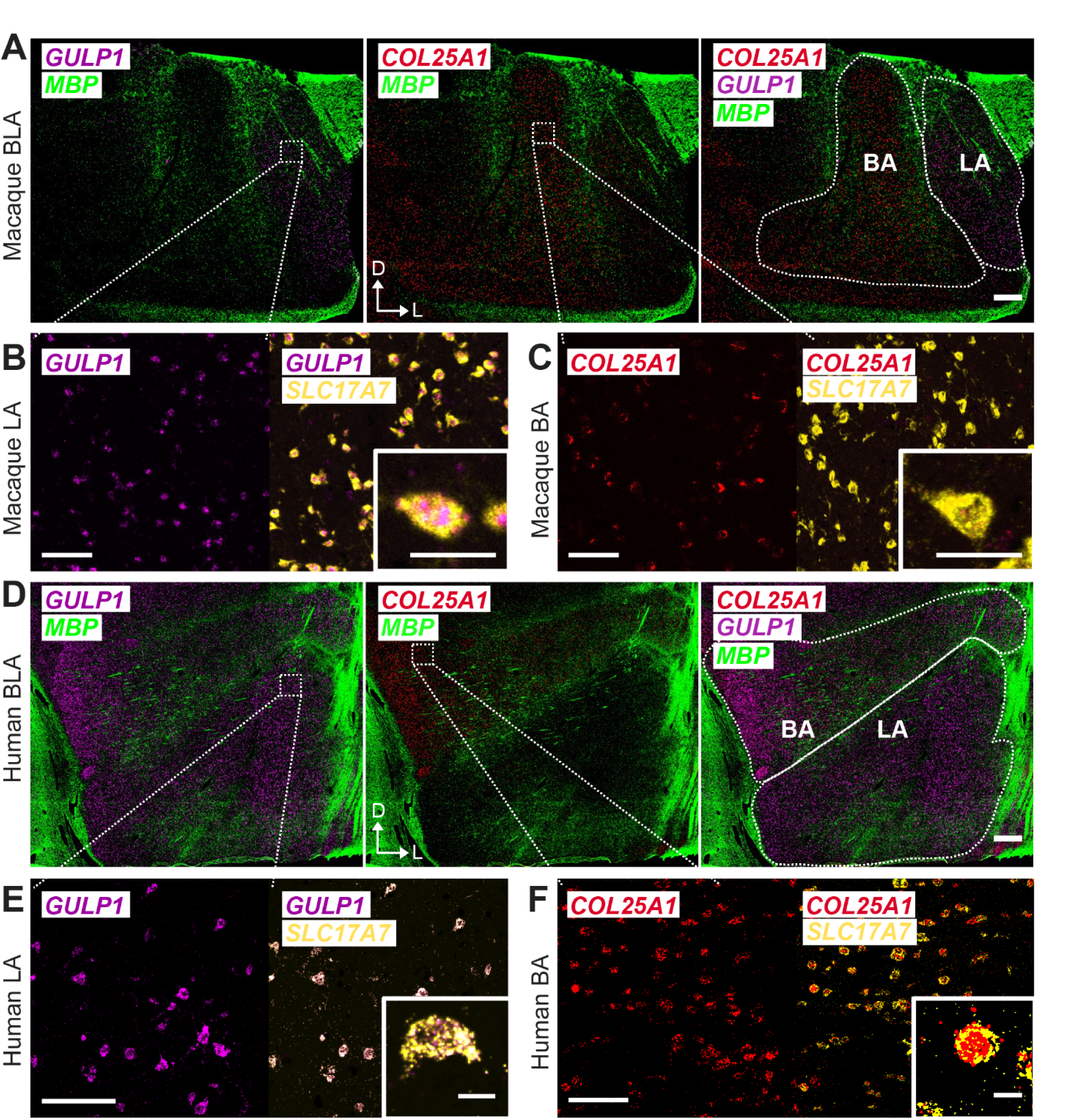
Marker gene expression for macaque and human lateral amygdala (LA) and basal amygdala (BA). 2X smFISH images of the basolateral region of the macaque (A) and human (D) brains illustrating expression of *GULP1* (magenta) in LA and *COL25A1* (red) in BA. *MBP* signal (green) represents white matter for anatomical landmarks. Dorval (D) and lateral (L) arrows are added for tissue directionality. White boxes represent approximate locations of zoomed in images. Scale bar 1000 µm. Zoomed in smFISH images illustrating co-expression of *GULP1* (magenta) and *SLC17A7* (yellow) within the 20X macaque (B) and 40X human (E) LA. Scale bar 100 µm. Inset illustrates a representative neuron co-expressing *GULP1* and *SLC17A7*. Scale bar of inset 10 µm. Zoomed in smFISH images illustrating co-expression of *COL25A1* (red) and *SLC17A7* (yellow) within the 20X macaque (C) and 40X human (F) BA. Scale bar 100 µm. Inset illustrates a representative neuron co-expressing *COL25A1* and *SLC17A7*. Scale bar of inset 10 µm.

We validated two additional markers of spatially-segregated excitatory neurons. We chose *PEX5L* as it was the top marker gene of the second excitatory neuron cluster specific to the basal nucleus, and *SATB1* as it was the second highest differentially expressed gene in lateral nucleus excitatory neurons in both macaques and humans (Fig. 4A and B). In macaques, there was clear spatial segregation of excitatory neurons expressing either *SATB1* or *PEX5L* (Fig. S17), with excitatory neurons expressing *SATB1* confined to the lateral nucleus, and neurons expressing *PEX5L* confined to the basal nucleus. Interestingly, the density of cells expressing *PEX5L* and *SATB1* was higher in the dorsal, magnocellular portions of the basal and lateral nuclei.

### Cryptic amygdala neurons with diencephalic origins are present in primates and humans

During global clustering of amygdalar snRNAseq profiles, we identified a population of *MEIS1^+^*/*PARD3B^+^* neurons that co-clustered with inhibitory rather than excitatory neurons in the UMAP embedding (middle panel, Fig. 1F). However, we manually annotated these neurons as excitatory based on enriched expression of *SLC17A6* coupled with a lack of canonical GABAergic neuron marker expression (e.g. *GAD1* or *GAD2*). Despite this expression profile, this neuron type was more closely associated with GABAergic rather than glutamatergic cell types. Cryptic glutamatergic neurons with transcriptional similarity to GABAergic neurons were also recently identified in marmosets and mice, and localized to the medial amygdala ^18^. The transcriptomic identity of these cryptic amygdala neurons in marmosets and mice suggests they cross from the diencephalon into the telencephalon early in development as their gene expression profiles overlap with those of GABAergic neurons in the hypothalamus ^18^. In marmosets, these cryptic amygdala neurons express the transcription factors *OTP* and *SIM1*. We also observed relatively high *OTP* and *SIM1* expression in the *MEIS1^+^/PARD3B^+^* cluster (Fig. S12). Moreover, in our macaque samples, *MEIS1^+^*/*PARD3B^+^* neurons came almost exclusively from samples of the central or dorsal basal nuclei (Fig. 4E), anatomical positions that unintentionally could include tissue from the medial amygdala (Fig 1A). These data implicate a similar pattern of development and migration for *MEIS1^+^*/*PARD3B^+^*neurons in human, baboon and macaque.

Interestingly, we also observed heightened *SLC17A6* expression coupled with lack of *GAD1*, *GAD2, or SLC32A1* expression in the *CARTPT^+^*/*CDH23^+^*cluster of inhibitory neurons. We classified this cluster of neurons as inhibitory because it broadly clustered with other GABAergic neuron types, and in macaques it was populated by neurons originating from the central nucleus. However, it is likely that these neurons are actually glutamatergic. For example, there are multiple CARPT^+^ glutamatergic neuron populations found in the mouse, especially in the thalamus ^71^. *CARTPT* is also widely expressed in the hypothalamus ^55^. Overall, this suggests that the *CARTPT^+^*/*CDH23^+^* is an additional class of cryptic amygdala glutamatergic neurons.

### Association of inhibitory cell types with neuropsychiatric disorders

Amygdala dysfunction is a feature of many neuropsychiatric disorders, particularly those associated with fear or anxiety, mood regulation, and addiction ^72–75^. We took two separate approaches to investigate potential association of discrete neuronal cell types with neuropsychiatric disorders (Fig. 7). First, we calculated statistical enrichment of genes previously identified to be downregulated in PTSD across all identified amygdala cell types, neuronal and non-neuronal ^27^, using only the human sn-RNAseq dataset. Specifically, we computed the ratio of genes that were statistically enriched in humans within a particular cell type to the total number of genes that were downregulated in patients diagnosed with PTSD. Using this metric, microglia showed the strongest association with patterns of decreased gene expression linked to PTSD diagnosis (Fig. 7A). This is consistent with prior reports that decreased expression of immune factors and microglia involvement are generally predictive of psychiatric diagnoses and served as a positive control. After correcting for multiple comparisons, all of the remaining cell types associated with statistical enrichment of genes downregulated in PTSD were classes of inhibitory neurons (Fig. 7A). This finding strengthens previous data suggesting that aberrant disinhibition of amygdala circuitry is causally linked to PTSD ^72,73,76^.

**Figure 7:**
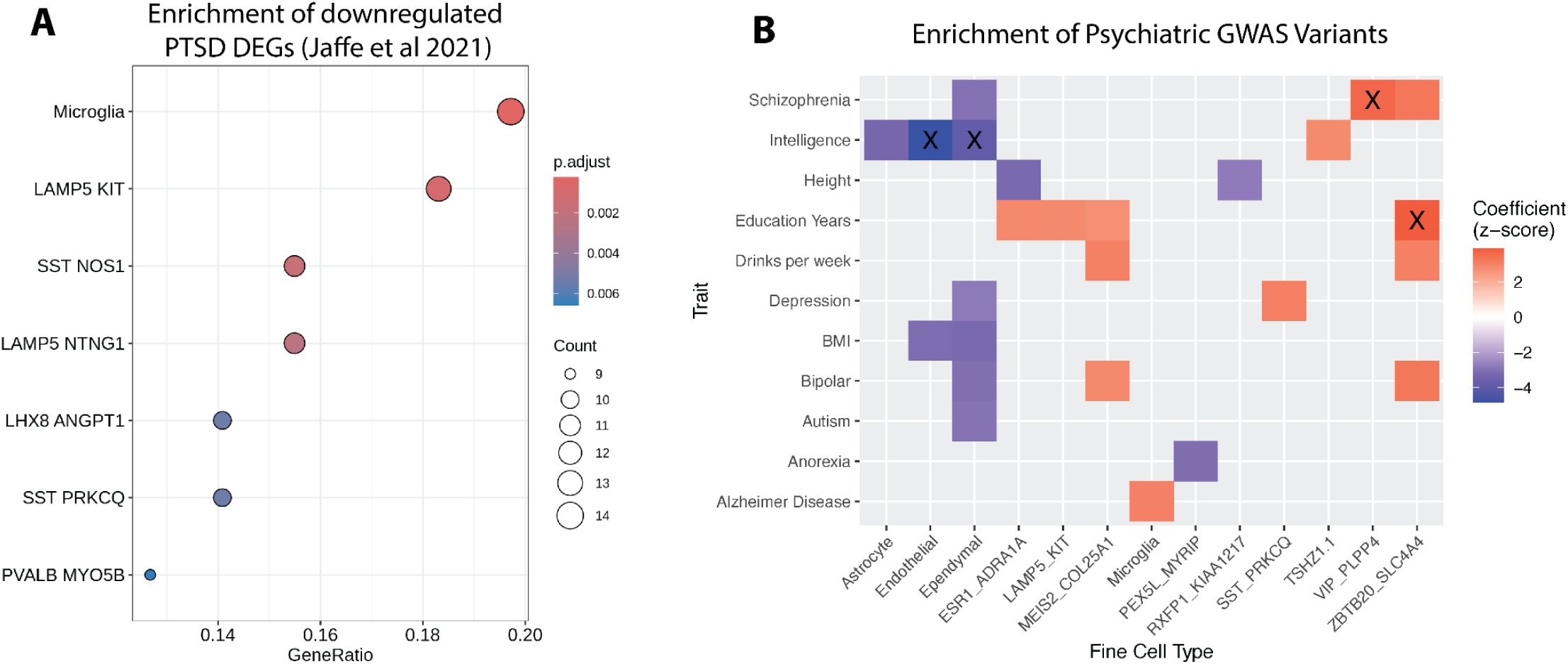
Association of amygdala cell types and subregions with psychiatric disorders. **(A)** Dot plot displaying cell type enrichment of downregulated differentially expressed genes (DEGs) in post-traumatic stress disorder (PTSD) from bulk RNAseq data in Jaffe et al (2021). Dots are colored by FDR-adjusted p-values and sized by the count of cell type marker genes found in the PTSD downregulated DEGs. **(B)** Heatmap showing Linkage Disequilibrium Score (LDSC) regression coefficient z-scores for heritability of different polygenic traits (y-axis) across cell types (x-axis). Only scores with FDR-adjusted *p* < 0.1 are colored. ‘X’ denotes FDR-adjusted *p* < 0.05. BMI, body mass index. Both analyses considered all cell types (i.e. non-neuronal, glutamatergic, and GABAergic neurons) but only significant associations are visualized.

The *LAMP5^+^*/*KIT^+^* and *LAMP5^+^*/*NTNG1^+^* neuron types are putative neurogliaform cells and it is notable that in macaques both types were more prevalent in the basal, accessory basal, and lateral nuclei than the central nucleus (Fig. 3D and Fig. S6). The *SST^+^*/*PRKCQ^+^* interneurons, like the *LAMP5^+^*interneurons, are more prevalent in the basal, accessory basal, and lateral nuclei than in the central nucleus (Fig. 3D). The *SST^+^*/*NOS1^+^*neurons are putative GABAergic projections and in macaques are more prevalent in the lateral than the basal nucleus (Fig. 3D). The *PVALB^+^*/*MYO5B^+^*interneurons were much more prevalent in the basal nucleus than any of the other nuclei (Fig. S6). The prevalence of *LHX8^+^*/*ANGPT1^+^*neurotensin releasing interneurons did not differ by amygdalar subregion.

Interestingly, *LAMP5^+^*/*KIT^+^* interneurons expressed corticotropin-releasing hormone (CRH; Fig. 3B), especially in humans and baboons. *CRH* and related genes are consistently downregulated in the amygdala of patients diagnosed with PTSD ^27^. *LAMP5^+^*/*KIT^+^* interneurons are also characterized by minimal expression of all but one of the genes encoding serotonin receptors (*HTR1E*; Fig 3I). *HTR1E* is specific to primates, but its function is largely unknown due to a lack of selective pharmacological tools and specific antibodies.

Next, we broadened the scope of our inquiry by using stratified-linkage disequilibrium score regression (S-LDSC; Fig. 7B). We first defined sets of differentially expressed genes for the cell types identified in our study. We then evaluated enrichment of genes linked to heritability of various traits or disorders among each cell type. Specifically, we used GWAS summary statistics for 21 traits. S-LDSC regression coefficients represent the contribution of a given cell type to the heritability of a given trait or disorder. In addition to the inclusion of neuropsychiatric disorders and traits related to these disorders, we also included height as a negative control and Alzheimer’s disease as a positive control. After correcting for multiple comparisons (FDR *p* < 0.05) we found two associations. Genes enriched in *VIP^+^/PLPP4^+^* interneurons were positively associated with genetic risk for schizophrenia, and genes enriched in *ZBTB20^+^/SLC4A4^+^* excitatory neurons found in the lateral nucleus were positively associated with education level. Using uncorrected statistical thresholds, there was an expected positive association between microglia gene expression and genetic risk for Alzheimer’s disease, as well as a previously described association between *SST+* interneurons and genetic risk for depression ^77^. We also found that excitatory neurons specific to the basal (*MEIS2^+^/COL25A1^+^*) or lateral nucleus (*ZBTB20^+^/SLC4A4^+^*) were positively associated with multiple traits: education level, bipolar disorder, and alcohol consumption. Finally, *ZBTB20^+^/SLC4A4^+^* excitatory neurons in the lateral nucleus were positively associated with schizophrenia. These associations are of interest given that alcohol use disorder, bipolar disorder, and schizophrenia are all characterized by alterations in emotion processing and linked to dysregulated amygdala function ^78–80^.

## DISCUSSION

Despite neuroanatomical evidence that the amygdala is a hub of interconnected nuclei that mediate specific motivated behaviors across species ^17,81–83^, it has remained challenging to understand structure-function relationships in the primate. Contributing to this challenge is a lack of knowledge about the molecular diversity of excitatory and inhibitory neurons within and across individual amygdalar nuclei. This has hindered primate neuroscientists’ ability to capitalize on available knowledge about cytological and histochemical features of the amygdala, as well as connectivity ^15^. Our approach took advantage of the spatial layout of the nonhuman primate amygdala to enable targeted sampling of five nuclear subdivisions allowing for spatially-resolved molecular profiling of excitatory and inhibitory neurons, which to date has only been available for rodents. We discovered that the primate amygdala contains at least 30 different populations of excitatory and inhibitory neurons, all of which are highly conserved across human and nonhuman primate species and many of which have distinct anatomical origins. Moreover, we demonstrated the utility of this anatomically-defined tissue dissection approach by showing how annotations of cell types in the nonhuman primate can infer the anatomical origins of human cell types. These results affirm the macaque and baboon as translationally-relevant experimental models for studying the function of constituent neuron types in humans.

We identified 12 different types of glutamatergic neurons in the primate amygdala that correspond to the pyramidal-like projection neurons identified in prior neuroanatomical studies ^15^, sometimes referred to as principal neurons ^43^. While differential assignment of functional roles for specific inhibitory neuron subtypes in amygdala are well-documented, heterogeneity amongst excitatory neurons is not assumed ^1^. Our results strongly argue against this perspective, which is also supported by the presence of multiple, molecularly-specialized glutamatergic neurons in the marmoset amygdala ^18^ and rodent amygdala ^4–8,84^.

The observed heterogeneity associates with multiple neurobiological features. A key feature being the nuclear subdivision to which an excitatory neuron cell body localizes. For example, 7 different types of excitatory neurons were found almost exclusively in either the accessory basal, basal, lateral, or central nucleus. While localization of the *ESR1^+^/ADRA1A^+^*neurons in the accessory basal nucleus and the *MEIS1^+^*/*PARD3B^+^*neurons in the central nucleus might have been predicted based on brain-wide expression patterns of *ESR1* ^68,69^ and *SIM1* ^18^, there is little neuroanatomical precedent for the other subdivision specific excitatory neuron types we identified. Prior morphological analyses of amygdala glutamatergic neurons across species have not convincingly identified more than one pyramidal-like cell type ^15,43^. Therefore, neuron type-specific differences in the morphology of the glutamatergic neuron types we identified are likely quantitative rather than qualitative. The known morphological differences between pyramidal-like excitatory neurons in the amygdala and pyramidal neurons in the cortex should provide insights into potential morphological attributes associated with different transcriptomic profiles ^18,19^. Amongst the excitatory neuron cell types identified, we observed differential expression of a number of transcription factors (*SATB2, MEIS1, MEIS2, TRPS1, ZBTB20*) implicated in the development of cortical glutamatergic neurons. Future experiments should utilize the molecular framework introduced in this atlas of the primate amygdala to identify morphological and connectivity features of glutamatergic amygdala neurons, including axonal project targets.

Another potential source of heterogeneity are differences in the sustained excitability of glutamatergic amygdala neurons. *In vitro*, pyramidal-like neurons in the amygdala exhibit three types of responses in response to a prolonged depolarizing input ^43,66,85^ when spike frequency adaptation is used as a quantitative metric of neuronal excitability. A small minority of neurons respond with a single spike at stimulus start, that is not accompanied by a sustained hyperpolarization. The majority of neurons spike repeatedly or in bursts at the start of the stimulus, and then exhibit spike frequency adaptation followed by an afterhyperpolarization, during which a subsequent response cannot be elicited. The remainder of neurons show minimal or no adaptation and fire until stimulus offset. Despite its utility as a quantitative metric, spike frequency adaptation has not been useful in identifying more than two types of excitatory neurons and it is not correlated with spatial location. This contrasts with *in vivo* responses to depolarizing inputs. Projection neurons in the basal amygdala of the cat respond with bursts of spikes, whereas projection neurons in the lateral nucleus respond with slow membrane potential oscillations ^86^. Similar differences in the excitability of neurons in the basal versus lateral nucleus are observed during neural recordings in awake, behaving macaques ^31^. Consistent with these findings we used gene ontology analyses of upregulated genes in glutamatergic neurons located in the lateral versus basal nucleus to identify enrichment of multiple GABA receptor subunits (*GABRA1* and *GABRB2*) and potassium voltage-gated ion channels (*KCNH5*, *KCBN2*, *KCNJ2*) in lateral nucleus excitatory neurons, that indicated neurotransmitter-gated ion channel activity regulates the postsynaptic membrane potential as a key biological process (Fig. S13).

The lateral and basal nuclei of the primate amygdala differ dramatically in terms of their connectivity and function ^10,11,15,31,87^. In comparison to the central nucleus, the basolateral complex has expanded considerably in primates to accommodate the expansion of the prefrontal cortex and binocular vision ^33,34,88^. Despite many causal experiments in nonhuman primates and humans implicating the amygdala in motivated behaviors ^9–12^ a largely unmet goal is direct comparisons of the effects on motivated behaviors after perturbing specific nuclei or cell types ^14^. For example, all prior studies that have manipulated amygdala function in nonhuman primates using chemogenetics reported that chemogenetic receptors were detectable in multiple nuclei ^89–91^. The identification and spatial validation of *GULP1*, *SATB1*, *COL25A1*, and *PEX5L* as genetic markers that target specific populations of excitatory neurons in nonhuman primates is a critical first step towards using molecular genetic tools to manipulate specific circuits within the primate amygdala. Importantly, two types of excitatory neurons that localized to either the basal (*MEIS2^+^*/*COL25A1^+^*) or lateral (*ZBTB20^+^*/*SLC4A4*) nucleus were associated with increased heritability of traits associated with psychiatric disorders. Manipulating these cell types in nonhuman primates may provide new opportunities for translational neuroscience with clinical relevance.

Prior attempts to annotate neurons from gross dissections of the human amygdala utilized marker genes identified in rodents ^20,21^. Yu and colleagues used this approach to identify several types of excitatory neurons that expressed genes which overlapped with the marker genes we identified, including *ESR1*, *COL25A1*, *SATB2*, and *ST8SIA2*. But they were unable to confidently assign anatomical origins for each cell type. For example, *ESR1* expression is highest in the posterior amygdala in rodent atlases, which corresponds to the accessory basal nucleus in the macaque amygdala ^15^. We confirm that *ESR1^+^*excitatory neurons are a highly conserved cell type across nonhuman primates and humans, and are localized to the accessory basal nucleus. Also, Yu et al. (2023) hypothesized that *SATB2^+^* and *LAMP5^+^* excitatory neurons were more prevalent in the lateral nucleus of the human amygdala. We found that *SATB2* expression was enriched in 3 specific types of excitatory neurons (*RXFP1^+^/KIAA127^+^, SLC17A8^+^/ST8SIA2^+^, and SATB2^+^/MPPED1^+^*) that represented a minority of the excitatory neurons analyzed. In macaques these cell types were more prevalent in the lateral nucleus, but were also found to an equivalent extent in the basal nucleus. In addition, we did find enriched expression of *LAMP5* in neurons specific to the lateral nucleus (*ZBTB20^+^*/*SLC4A4)*, but we also detected enriched expression in the *MEIS2^+^*/*COL25A1^+^*and *ESR1^+^/ADRA1A^+^* neurons with anatomical origins in the basal and accessory basal nucleus. Finally, many of the marker genes previously found to dissociate neuron types in the basal versus lateral nucleus of the rodent amygdala did not discriminate the anatomical origins of neurons in the primate amygdala (Fig. S14). The ability to detect similar neuron types using marker genes from rodent atlases, but the inability to replicate the spatial heterogeneity we have identified in the present atlas, emphasizes the continuing need for constructing spatially-resolved transcriptomic atlases in multiple species of primates. Targeted tissue dissections are one approach to maintaining spatial information as high-throughput viral connectomic approaches ^92,93^ are developed for use in nonhuman primates.

One advantage of our sampling approach was the ability to compile molecular profiles of all GABAergic neurons across the primate amygdala. Prior neuroanatomical studies have examined specific classes of inhibitory neurons, but in a piecemeal fashion ^15^. A standout result from the construction of this atlas is the large dissociation in the ratio of GABAergic to glutamatergic neurons in the central nucleus versus other subdivisions. On balance the central nucleus consists mainly of GABAergic neurons, whereas the opposite is true for the accessory basal, lateral, and basal nuclei. This profile for the central nucleus fits with evidence that the central nucleus of the amygdala shares developmental origins with the striatum ^94^

The central nucleus was the only subdivision to uniquely contain specific types of GABAergic neurons, particularly interneurons expressing *PKRCD*. We identified two types of *PKRCD^+^* interneurons based on their expression of either the D1 or D2 dopamine receptor. Prior investigations of *PKRCD^+^*interneurons in rodents focused on D2-mediated effects on aversive learning and fear expression because of its abundant expression in the central nucleus ^37,95^. In rodents, threatening stimuli activate SST-containing interneurons that in turn inhibit *PKRCD^+^* interneurons leading to the disinhibition of brainstem circuits mediating autonomic and reflex responses. Likewise direct activation of *PKRCD^+^* interneurons has anxiolytic effects ^1^. Consistent with direct manipulations of this neuron type in rodents, individual differences in *PKRCD* expression in rhesus macaques is moderately associated with behavioral measures of anxious temperament and a subset of *PKRCD* interneurons innervated by SST-containing interneurons project to the bed nucleus of the stria terminalis (BNST) ^38^. The primate BNST is characterized by dense expression of D1 and D2 receptors ^96^ so we cannot infer which of the two *PKRCD^+^* neuron types are mediating a potential relationship with anxious temperament. Also, despite claims regarding the role of the central nucleus in anxious temperament ^97^ we found that many of the genes associated with anxious temperament in prior studies (e.g. *SS18*, *DNMT3A, NTRK3, IRS2*) ^38,98^ were either expressed in multiple glutamatergic and GABAergic neuron types located outside the central nucleus or showed ambient levels of gene expression (e.g. *SLC18A2, KIAA1009)*. Considering the similar developmental origins of the striatum and central nucleus it is interesting to consider if differences in D1 and D2 receptor expression amongst *PKRCD^+^* interneurons plays a similar role in shaping appetitive and aversive responding, as in the striatum ^99^. Modulation of D1 and D2 receptor function in the central nucleus has mixed effects on fear learning ^64^ and extinction ^95^. The complexities of dopamine receptor expression within the central nucleus and *PKRCD^+^*interneuron subtypes warrant further investigation, especially in nonhuman primates.

The intercalated cell masses are similarly understudied in primates relative to rodents, despite the key role of the ITCs in amygdala function. For example, there have been no functional characterizations of ITCs in primates, despite neuroanatomical validations of *FOXP2* and *TSHZ1* as conserved markers of ITC neurons ^18,25,26^. We identified two distinct clusters of *TSHZ1^+^/FOXP2^+^*interneurons that were highly conserved across nonhuman primates and humans and that differed in their expression of the D3 dopamine receptor and 5-HT7 serotonin receptor (Fig. 3E and G). Dopamine affects the plasticity of ITC neurons that form synapses with excitatory projection neurons from the lateral nucleus ^64^, but serotonergic modulation of ITC neuron plasticity and function is understudied. The two ITC neuron types also differed in expression of genes that regulate striatal projection neuron development, suggesting that the D3-enriched population was spiny and the HTR7-enriched population was aspiny, confirming a prior morphological analysis of ITC neurons ^26^. In the marmoset, *FOXP2^+^* GABAergic amygdala neurons and striatal projection neurons have similar gene expression profiles ^56^, but their anatomical origins remain unknown. In macaques the larger cluster of *TSHZ1^+^/FOXP2^+^* spiny neurons we identified mostly came from tissue samples of the central nucleus, whereas the smaller cluster of *TSHZ1^+^/FOXP2^+^* aspiny neurons were equally distributed across all nuclear subdivisions. These spatial patterns are consistent with those from prior neuroanatomical studies of the ITC in nonhuman primates ^26^. Because direct dissection of the ITC is impossible without molecular validation, future experiments can capitalize on our molecular profiling to utilize spatial transcriptomics to better map the distribution of these two neuron types, especially in humans. Such efforts will be instrumental for understanding how ITC neurons contribute to amygdala function in primates.

## Conclusion

Transcriptomic profiling makes it clear that heterogeneity amongst glutamatergic and GABAergic amygdala neurons is a rule rather than an exception. In conjunction with prior neuroanatomical and electrophysiological studies, our data demonstrate that heterogeneity in molecular profiles originates based on where a neuron is located within the amygdaloid complex and what neurotransmitters, hormones, and peptides affect its function. A necessary next step is further relating this heterogeneity to quantitative differences in neuronal morphology, connectivity, and function. Going forward, conceptual and computational models of amygdala function will need to account for this heterogeneity to accurately predict amygdala-dependent changes in behavior that are relevant for mental health disorders. The identification of conserved genetic markers associated with excitatory neuron populations specific to the basal or lateral nucleus in nonhuman primates overcomes a major hurdle to building such models. The transcriptomic profiles described here represent a first step to discovering genetic elements associated with molecular markers that can be used to build tools for selective manipulation of individual amygdala nuclei. Such studies will be critical to determine the relevance of the nuclei-specific cell types for amygdala related behaviors and disease.

## MATERIALS AND METHODS

### Post-mortem nonhuman primate tissue samples

Fresh, unfixed, and flash frozen amygdala tissue punches were acquired from 5 rhesus macaques and 2 female olive baboons (Supplementary Table 1). All animals were paired housed on a 12-hour on/12-hour off lighting schedule with ad libitum access to food and water. Animals were fed standard primate chow twice daily and provided fruit and vegetable enrichment daily. Macaques were observed by trained veterinary technicians daily in their home cages. The Institutional Animal Care and Use Committee and the Institutional Biosafety Committee at the ONPRC and Oregon Health and Science University (OHSU) approved all experimental procedures, and all of the guidelines specified in the National Institutes of Health Guide for the Care and Use of Laboratory Animals (National Research Council, 2011) were strictly followed.

Necropsies and tissue collections were performed as previously described ^100,101^. Animals were sedated with ketamine (10 mg/kg) and then deeply anesthetized with sodium pentobarbital followed by exsanguination. The brain and spinal cord were perfused through the ascending carotid artery with 1 L of 0.9% ice cold saline. The brain was then removed from the skull (< 30 minutes post-mortem), deposited into an ice-cold bath of saline for transport, and placed into an ice-cold, steel brain matrix (Electron Microscopy Sciences). A custom 3-D printed brain matrix was used to section the baboon brains. Each brain was positioned in the brain matrix with the ventral surface facing up. The anterior medial temporal sulcus, posterior to the temporal pole and anterior to the rhinal sulcus was identified on the ventral surface of the temporal lobe. A carbon steel knife blade (Thomas Scientific) was inserted into the slot in the brain matrix that was most closely aligned and orthogonal to the beginning of the anterior medial temporal sulcus. Additional knife blades were inserted anterior and posterior to the first knife blade in 2 mm increments. Depending on the size of the brain this resulted in either 2 to 3 brain slabs that encompassed the anterior posterior extent of the amygdala. The resulting bilateral brain slabs were then removed from the brain matrix and laid out flat in sterile petri dishes treated with RNAase-X. The petri dishes rested on an aluminum plate secured to a chamber filled with dry ice. The temperature of the plate was monitored every 5 minutes with an infrared thermometer to maintain a temperature of -30 to -15 C for up to two hours while tissue punches were acquired and flash frozen.

Brain slabs in which the amygdala was clearly visible below the anterior commissure or globus pallidus on both sides of the slab were identified and 1.0-2.5 mm in diameter tissue punches were taken through the full width of the slab. For posterior slabs in which the amygdala appeared on the anterior face of the slab and the hippocampus appeared on the posterior face of the slab, tissue punches were collected, carefully removed from the biopsy needle, and the rostral aspect of the punch was retained for nuclei isolation and snRNAseq. Tissue punches were checked to ensure they did not contain noticeable quantities of white matter and then inserted into DNAase and RNAase-free 1.5 mm LoBind (Eppendorf Protein LoBind Tube, Cat #22431102) microcentrifuge tubes that were inserted into pulverized dry ice. Ethanol was poured onto the dry ice to flash freeze the tissue punches. All tissue punches were acquired less than 90 minutes postmortem. The punched brain slabs were then post-fixed in 4% paraformaldehyde for 48 hours, cryoprotected in 30% sucrose, and then sectioned in 40 µm sections for histological analyses.

### Postmortem human tissue samples

Post-mortem human brain tissue was obtained at the time of autopsy with informed consent from the legal next of kin, through the Office of the Chief Medical Examiner of the State of Maryland, under the Maryland Department of Health IRB protocol #12–24, the Departments of Pathology at Western Michigan University Homer Stryker MD School of Medicine, and at the University of North Dakota School of Medicine and Health Sciences, and through Gift of Life Michigan, all under WCG IRB protocol #20111080. Demographics for the 5 donors are listed in Supplementary Table 2. Details of tissue acquisition, handling, processing, dissection, clinical characterization, diagnoses, neuropathological examinations, and quality control measures have been described previously ^102^. Frozen coronal brain slabs containing the amygdala at the level of clearly visible caudate nucleus and putamen (separated by the internal capsule), globus pallidus external and internal segments, fornix, and the anterior commissure were selected for dissections. Tissue blocks (approximately 10mm x 20mm) containing the amygdala were dissected under visual guidance with a hand-held dental drill. Blocks were stored in sealed cryogenic bags at -80°C until cryosectioning and between tissue collection rounds. At the time of cryosectioning, tissue blocks were acclimated to the cryostat (Leica CM3050) at -14°C, mounted onto chucks, ∼50 µm of tissue was trimmed from the block to achieve a flat surface, and several 10 µm sections were collected for quality control (RNAscope, H&E). After identification of the boundaries of the basolateral complex (BLA), blocks were again acclimated to the cryostat, mounted onto chucks, and scored with a razor to isolate the BLA. 100 µm sections of the BLA were collected in pre-chilled 2 mL DNAase and RNAase-free microcentrifuge tubes (Eppendorf Protein LoBind Tube, Cat #22431102) for a total of 70-100 mg of tissue and stored at -80°C until nuclear isolation.

### Anatomical validations and quality control of human tissue blocks (H&E and RNAscope)

Prior to collecting sections for snRNAseq experiments, tissue blocks were cut and 10 µm sections were collected to complete two quality control steps: 1) H&E staining to assess gross neuroanatomical structure of the block, and 2) single molecule fluorescence *in situ* hybridization (fluorescence multiplex RNAscope). H&E staining and RNAscope were performed according to manufacturer’s instructions and images were acquired using an Aperio CS2 slide scanner (Leica) or a NikonAXR (Nikon Instruments), respectively. For RNAscope, probes for established marker genes (ACD Bio) were used to identify white matter and subnuclei of the amygdala, including *MBP* (Cat # 411051), *SLC17A7* (Cat # 415611), *COL25A1* (Cat # 1187021)*, GULP1* (Cat #1095761), *PEX5L* (Cat # 1185381), *SATB1* (Cat # 454621).

### RNAscope single molecule fluorescent in *situ* hybridization (smFISH)

Fresh frozen amygdala from rhesus macaque donor and human donor were sectioned at 10 µm and stored at -80 °C. *In situ* hybridization assay was conducted using RNAscope Multiplex Fluorescent Reagent Kit v2 (Cat # 323100, ACD, Hayward, California) following manufacturer’s protocol. In summary, the tissue sections were fixed in 10% Neutral Buffered Formalin (NBF) solution (Ct # HT501128-4L, Sigma-Aldrich, St. Louis, Missouri) for 30 minutes at room temperature. Sections were dehydrated with serial ethanol washes, pretreated with hydrogen peroxide for 10 minutes at room temperature, and treated with protease IV for 30 minutes. Four different probe combinations were used at one time to identify marker gene expression in the BLA with the following probes: Hs-SLC17A7 (Cat #415611-C4, ACD, Hayward, California), Hs-COL52A1 (Cat # 1187021-C2, ACD, Hayward, California), Hs-GULP1 (Cat # 1095761-C3, ACD, Hayward, California), Hs-MBP (Cat # 411051-C4, ACD, California Hayward), Hs-PEX5L (Cat # 1185381-C2, ACD, California Hayward), Hs-SATB1 (Cat # 454621, ACD, Hayward, California), Mmu-GULP1-C3 (Cat # 1568911-C3, ACD, Hayward, California), Mmu-MBP-C4 (Cat # 1006431-C4, ACD, Hayward, California). After labeling the probes, sections were stored overnight in 4x SSC buffer. Following amplification steps (AMP1–3), the probes were fluorescently labeled using opal dyes (Perkin Elmer, Waltham, MA; diluted 1:500) and counterstained with DAPI (4′,6-diamidino-2-phenylindole) to mark the nucleus. Sections were imaged using 2X - 40X objectives on NikonAXR (Nikon Instruments).

### Nonhuman primate nuclei isolation and snRNAseq

Flash-frozen tissue punches collected from the different subdivisions of the amygdala of macaques (4 subdivisions) and baboons (2 subdivisions) were dissociated into individual nuclei following manufacturer’s instructions (protocol CG000366, 10X Genomics). Briefly, a chilled lysis buffer was added to a chilled glass dounce and the cryosections were transferred while frozen. Sections were homogenized using 5 or 15 strokes, respectively, with both loose and tight-fit pre-chilled pestles. Then, the lysis buffer was neutralized with a resuspension buffer. The homogenate was strained through a 70 µm cell strainer, followed by a 40 µm cell strainer. Nuclei suspension were centrifuged in a 29-50% OptiPrep gradient (Sigma-Aldrich) at 3000 g for 20 min at 4C to separate intact nuclei from debris. The pelleted nuclei was resuspended and centrifuged at 500 g at 4°C for five minutes, and the supernatant was removed for a total of three times. Nuclei were inspected in a counting chamber for intact, bright, nongranular cell morphologies, indicating high viability and successful debris removal. Approximately 10,000 single nuclei were captured for each sample in a single channel on the 10X Chromium controller, and snRNAseq libraries (Chromium Next GEM Single Cell 3’ kit v3.1) were generated following the manufacturer’s instructions. Libraries were sequenced to an average depth of 20,000 read pairs per nucleus, in a NovaSeq 6000 at the Oregon Health and Science University Massively Parallel Sequencing Shared Resource according to 10X Genomics’ specifications. All samples were processed and sequenced at the same time to avoid batch effects. For the baboon samples, nuclei were extracted from flash-frozen tissue and enriched for intact, single nuclei from neurons using fluorescence activated nuclei sorting (FANS) based on NEUN immuno-labeling (using Rabbit-anti NEUN monoclonal anti-body directly conjugated to Alex Fluor 594; Cell Signaling Technology, Cat #: 90171S) and DAPI staining using a slightly modified version of a previously published protocol ^103^. FANS parameters - including NEUN gating thresholds and nuclei recovery rates were established for each experiment using un-stained control nuclei and approximately 20,000 NEUN+/DAPI+ nuclei were loaded into each Next GEM reaction. Library generation and sequencing were performed as described for macaque samples.

### Human nuclei isolation and snRNAseq

Using 100 μm cryosections collected from each donor, we conducted snRNAseq using 10x Genomics Chromium Single Cell Gene Expression 3’ kit v3.1. Approximately 70-100 mg of tissue was collected from each donor, placed in a pre-chilled 2mL microcentrifuge tube (Eppendorf Protein LoBind Tube, Cat #22431102), and stored at -80C until the time of experiment. Nuclei preparations were conducted according to the 10x Genomics customer-developed “Frankenstein” nuclei isolation protocol, as previously described in ^21^, with modifications designed to optimize the protocol for use with cryosections. Briefly, chilled EZ lysis buffer (MilliporeSigma #NUC101) was added to the LoBind microcentrifuge tube containing cryosections, the tissue was fragmented by pipette mixing, this lysate was transferred to a chilled glass dounce, the tube was rinsed with additional EZ lysis buffer, which was added to the respective dounce. Sections were homogenized using 10-20 strokes with both loose and tight-fit pre-chilled pestles, and the homogenate was strained through a 70 µm cell strainer. After lysis, samples were centrifuged at 500g at 4°C for five minutes, supernatant was removed, and the pellet was resuspended in EZ lysis buffer and re-centrifuged. The supernatant was removed, and wash/resuspension buffer (PBS containing 0.5% BSA (Bovine Serum Albumin, Jackson ImmunoResearch #001-000-162) was added to the pellet. Upon resuspension, the samples were spun again and this wash process with wash/resuspension buffer was completed three times. Nuclei were labeled with Alexa Fluor 488-conjugated anti-NeuN (MilliporeSigma, Cat. # MAB377X) diluted 1:1000 in a nuclei stain buffer (1x PBS, 3% BSA, 0.2U/μL RNase Inhibitor), by incubating at 4°C with continuous rotation for 1 hour. Proceeding NeuN labeling, nuclei were washed once in stain buffer, centrifuged, and resuspended in wash/resuspension buffer. Nuclei were labeled with propidium iodide (PI) at 1:500 in wash/resuspension buffer and subsequently filtered through a 35μm cell strainer. Fluorescent activated nuclear sorting (FANS) was performed using a Bio-Rad S3e Cell Sorter at the Lieber Institute for Brain Development. Gating criteria were selected for whole, singlet nuclei (by forward/side scatter), G0/G1 nuclei (by PI fluorescence), and neuronal nuclei (by Alexa Fluor 488 fluorescence). 9,000 nuclei were sorted into a tube based on both PI+ and NeuN+ fluorescence to facilitate enrichment of neurons, which resulted in 81.9% neurons across all samples. Samples were collected over two rounds, each including 2 or 3 donors. All samples were sorted into reverse transcription reagent master mix from the 10x Genomics Single Cell 3′ Reagents kit (without enzyme). Reverse transcription enzyme and water were added to bring the reaction to full volume after the sort. cDNA synthesis and subsequent library generation was performed according to the manufacturer’s instructions for the Chromium Next GEM Single Cell 3’ v3.1 (dual-index) kit (CG000315, revision E, 10x Genomics). Samples were sequenced on a Nova-seq 6000 (Illumina) at the Johns Hopkins University Single Cell and Transcriptomics Sequencing Core.

### Processing of raw snRNA-seq data

All FASTQ files for snRNA-seq libraries were aligned using 10x Genomics software, cellranger count (v7.0.0). Libraries from the human donors were aligned to the Human genome reference (GRCh38/Hg38, Ensembl release 98), rhesus macaque samples were aligned to the macaque (*Macaca mulatta*) genome reference (Mmul_10, Ensembl release 110), and baboon samples were aligned to the Olive baboon (*Papio anubis*) genome reference (Panubis1.0, NCBI release 104). Feature-barcode files were analyzed in R v4.3.1 within the Bioconductor framework (v3.17), unless otherwise stated. Empty droplets were identified and removed using the emptyDrops() function from the DropletUtils package using a data-driven threshold. To identify droplets containing more than one nuclei (i.e., doublets), sample specific doublet scores were calculated using computeDoubletDensity() from snDblFinder v1.14.0 with the top 2000 highly variable genes. Droplets with a score greater than or equal to 2.75 were excluded from downstream analyses.

For human and baboon samples, quality control was performed by computing sample-wise median absolute deviation (MAD) thresholds for total number of unique molecular identifiers (UMI), number of unique detected genes, and the proportion of reads mapping to mitochondria. Individual cells were considered low quality if they exceeded less than -3 MADs for total UMI and unique genes and/or greater than +3 MADs for mitochondrial ratio, per sample. Total UMI and unique gene QC metrics for macaque samples displayed bimodal distributions due to the inclusion of both neuronal and non-neuronal cell types, resulting in no nuclei being excluded using the 3 MAD thresholds. Thus, minimum thresholds of 600 UMI and 500 unique genes were used instead. These preprocessing steps resulted in a final total of 21,020 human, 104,797 macaque, and 46,111 baboon nuclei.

To allow for cross-species comparisons, genes were subset to include only one-to-one orthologs using the convert_orthologs function from the orthogene package using the gprofiler method. For non-human primate data, input_species was set to either “macaque” or “baboon” and the output_species variable was set to “human”, and the strategy for handling non one-to-one matches was set to “drop_both_species”. The intersection was then taken of the one-to-one orthologs between the three species. After filtering out mitochondrial and lowly expressed genes, this resulted in a total of 13,842 common genes across human and non-human primate species.

### Feature selection, cross-species integration, and fine resolution clustering

Feature section, dimensionality reduction, cross-species integration, and clustering were all carried out using Seurat v4 workflows. The combined data were first split by subjects and species. All data were then separately normalized using the Normalizedata function, the top 2000 highly variable genes were selected using the FindVariableFeatures function, and the data were centered and scaled using the ScaleData function. Features used for integration were then selected using the SelectIntegrationFeatures function. These features were then centered and scaled for each data split before performing principal component analysis (PCA). Data integration was performed using the IntegrateData function using anchors determined from the FindIntegrationAnchors function. Default parameters were used for all functions.

After cross-species integration, data were re-scaled and PCA was run on the integrated data. Data were then clustered (FindClusters function) on the top 30 principle components using the Leiden algorithm and a resolution of 0.5. Data were visualized with the uniform manifold approximation and projection (UMAP) dimensionality reduction technique via Seurat’s RunUMAP function with the top 30 principle components. Default parameters were used for all functions, unless otherwise specified. These initial clusters were then grouped based on canonical marker gene expression into either excitatory neurons (*SNAP25*+,*SLC17A7*+, *SLC17A6*+), inhibitory neurons (*SNAP25*+, *GAD1*+, *GAD2*+), or non-neuronal cell-types (*SNAP25*-). Inhibitory neurons were also identified based on canonical marker genes such as *SST, PVALB, CARTPT, VIP, CCK, LAMP5, PRKCD, TSHZ1, NTS,* and *PENK.* To obtain fine resolution cell-type clusters, both excitatory and inhibitory neuron-types were sub-clustered by subsetting the data to exclusively excitatory or inhibitory neurons before rerunning the above Seurat integration and cluster workflows. Subcluster quality was assessed with the pairwise modularity ratio using the pairwiseModularity function from the bluster package, which determined how separated each cluster is from one another. Any clusters that showed poor separation (i.e., low modularity score) were collapsed to a single cluster.

### Cell type-specific marker genes, annotations, and visualizations

For excitatory neurons, fine-resolution cell type annotations were performed automatically using cluster-specific marker genes detected by findMarkers_1vAll function from the DeconvoBuddies package using the log-normalized counts with ∼species used as a blocking factor. Annotation names were then assigned based on the top two marker genes for each cell type as *Gene1_Gene2*. The same method was used for annotating inhibitory neurons, except canonical marker genes based on established classes of GABAergic cell types (*PVALB, SST, VIP, CCK, PRKCD, NTS, CARTPT, LAMP5,* and *TSHZ1*) were listed in place of *Gene1*. Non-neuronal cell types were also broadly labeled based on canonical gene expression for astrocytes (*GFAP*, *SLC1A1*), OPCs (*CD9*), oligodendrocytes (*MBP*), ependyma (*FOXJ1*), endothelial (*RGS5*), and microglia (*TMEM119*).

Heatmaps visualizing marker gene expression for excitatory, inhibitory, and non-neuronal cell types were all generated using the ComplexHeatmap package. For this, the mean log-normalized counts were aggregated across cells using the aggregateAcrossCells function from the scuttle package. Species proportions were calculated as the proportion of nuclei sampled from each species within each fine cell type. Because precise punches of all regions (LA, BA, aBA, and CeA) were only collected within macaques, the subregion proportions were calculated as the proportion of nuclei sampled from each subregion only within the macaque dataset.

### Putative subregion superclusters

Excitatory cell type clusters found to be enriched in lateral (*GULP1_TRHDE*, *ZBTB20_SLCA4*), basal (*PEX5L_MYRIP*, *MEIS2_COL25A1*), or accessory basal (*ESR1_ADRA1A*, *GRIK3_TNS3)* samples from NHPs were classified as putative LA and BA excitatory clusters in all species. These classifications were used in subsequent analyses for comparisons across amygdala subregions.

### Cross-species comparisons of fine cell types using MetaNeighbor

We used the MetaNeighbor package to assess cell type conservation across species using the unsupervised approach ^104^. This method evaluates the similarity between cell types across different datasets (species) by using one cell type as a training set and another as a test set, averaging the AUROC scores across these comparisons to create a matrix that reflects the degree of similarity between cell types across species. An AUROC score of 0.5 indicates random classification, whereas higher scores indicate high accuracy in predicting cell types across species. This allows us to determine the degree of conservation across cell types. Importantly, MetaNeighbor has been shown to be robust to class imbalances ^105^. Highly variable genes (selected with MetaNeighbor’s variableGenes function) were used as input features to the MetaNeighborUS function with the parameters study_id=”species”, cell_type=”fine_celltype”, fast_version=TRUE, one_vs_best=TRUE, and symmettric_output=FALSE. Heatmaps and boxplots were generated from the resulting AUROC matrix using the ComplexHeatmap and ggplot packages, respectively.

### Pseudobulk differential expression analysis

Novel marker genes for amygdala subregion were identified using pseudobulk differential expression analysis. The mRNA counts of excitatory neurons from independent lateral (n=15; 12 macaque and 3 baboon) and basal (n=17; 13 macaque and 4 baboon) samples were aggregated (i.e., pseudobulked) using the aggregateAcrossCells function from the scuttle package. Lowly expressed genes were filtered out using the filterByExpr function from the edgeR package with the group variable set to Subregion.

Pseudobulk DE analysis was carried out using the DESeq2 pipeline. In short, a DESeqDataSet was built using the DESeqDataSetFromMatrix with the raw aggregated counts. A design matrix of ∼species+subregion was created to analyze differences in subregions while controlling for differences across nonhuman primate species. To ensure no samples were obvious outliers, PCA was performed on the regularized log-counts (DESeq2’s rlog function) and the first two principal components were visualized using the autoplot function from the ggfortify package. After ensuring all samples were of high quality, default DE analysis was performed via the DESeq function, lateral and basal samples were contrasted using the results function, and the calculated log-fold changes (logFC) were shrunk using the lfcShrink. Only genes with *p*-values < .05 after correcting for the false discovery rate (FDR) were considered as candidate markers. Candidate marker genes were then filtered to only include highly expressed genes (mean counts >3000) and relatively large log FC (absolute_value(LFC) > 1). Volcano plots were generated using ggplot. To extend these findings to humans, using identical methods as above, putative human lateral and basal excitatory neuron clusters were pseudobulked within samples (n=5) and the expression of rlog-normalized counts of top candidate marker genes were visualized to confirm lateral and basal excitatory marker genes in humans using custom boxplots generated by ggplot. Identical methods were used for pseudobulk DE of intercalated cell type clusters (TSZH1.1 vs TSZH1.2) and PRKCD+ cell types (PRKCD_DRD1 vs PRKCD_DRD2).

### Correlation of marker gene t-statistics across species

To determine the cross-species conservation of lateral and basal marker genes, we ran a correlation analysis of the standard logFC of of marker genes associated with amygdala subregions. Excitatory neurons were first subset to just include nuclei within the putative lateral, basal, and accessory basal cell type clusters. Marker genes for the putative subregions were then found within each species using the findMarkers_1vAll function from the DeconvoBuddies package, while including Sample as a blocking factor. We then calculated the Pearson correlation coefficient (cor function with default parameters from base R) to compare the subregion marker gene logFC between species.

### Gene Ontology (GO) over-representation analysis

GO over-representation analysis of subregion pseudobulk differentially expressed genes (DEGs) was carried out via the compareCluster function using the clusterProfiler package. In brief, over-representation analysis was performed using parameter fun=enrichGO using the human genome annotations (org.Hs.eg.db) for each Cellular Components, Biological Processes, and Molecular Function ontologies. The Benjamin-Hochberg procedure was used for p-value correction and a q-value cutoff of 0.05 was used. To reduce redundancy in GO terms, we use the simplify function on output GO results with a cutoff of 0.7. The top 5 categories within each cell type or subregion per ontology were then visualized were then visualized use the dotplot function from clusterProfiler.

### Cell type-specific enrichment analysis (CSEA)

To determine which cell types may be implicated in psychiatric disease, we performed CSEA on PTSD and major depressive disorder (MDD) DEGs from a previously published bulk RNAseq study ^27^ using the enricher function from the clusterProfiler package. To allow for direct one-to-one comparison to the human from Jaffe and colleagues., the data were first subset to only include nuclei from human samples (*n* = 20,655 nuclei) and the gene set was expanded to include all sampled counts in the analyses (*n* = 21,844 genes) rather than only including one-to-one orthologs. After renormalizing the human dataset, the top 100 marker genes based on standard logFC were found for each cell type using the findMarker_1vAll function from the DeconvoBuddies R/Bioconductor package. DEGs were obtained from Jaffe et al. which included both DEGs associated with PTSD and MDD within the basolateral complex. Because only a small number of genes had an FDR-adjusted p-value < 0.05, genes with unadjusted p-values < 0.05 were included in the analysis and then classified by their logFC as up- or downregulated. For each condition, over-representation analysis was performed using the enricher function from the clusterProfiler package. No significant hits were found for enrichment of specific cell type markers in up-regulated genes for either PTSD or MDD gene sets. Enrichment of cell type markers in down-regulated were visualized using dot plots displaying gene ratios and adjusted p-values.

### Stratified linkage disequilibrium score regression (S-LDSC)

Prior to S-LDSC, we defined gene sets for each set of fine cell type classes using human data. In short, fine cell types were first filtered to exclude human cell types with less than 70 total cells before being pseudobulked using the aggregateAcrossCells function using default parameters. Genes were then filtered to retain only protein-coding genes that were expressed above background levels in a sufficient number of cells using the filterByExpr function from the edgeR package. This resulted in a total of 15,128 genes. The resulting pseudobulked dataset was then normalized by first calculating normalization factors (calcNormFactors function) and then normalizing to counts per million (cpm function) using the edgeR package.

Genes sets for use in S-LDSC were created by calculating the top 10% most highly expressed genes within each cell type. To further normalize gene expression, the expression of each gene was divided by the gene’s total expression across all cell types such that each gene’s expression values summed to 1. We then calculated the 90th percentile threshold for each cell type to define the top 10% highly expressed genes per cell type. We then added a 100Kb window up and downstream of the transcribed region of each gene in the gene sets to construct a genome annotation for each unique cell type label.

S-LDSC analysis was performed as previously described ^106^. In brief, we conducted stratified LD score regression (S-LDSC) to assess the heritability of brain-related traits across the gene sets associated with each human cell type. To determine whether the observed enrichments were specific to brain-related traits, we included human height and type 2 diabetes as negative control traits. GWAS summary statistics for all 21 traits were obtained from the respective studies ^107–115^. We performed S-LDSC using the baseline LD model v2.2, which includes 97 annotations to account for linkage disequilibrium with other functional genomic annotations, per the guidelines from the LDSC resource website (https://alkesgroup.broadinstitute.org/LDSCORE). We used SNPs from the HapMap Project Phase 3 as the regression SNPs and SNPs from the 1000 Genomes Project (European ancestry samples) as the reference SNPs, both sourced from the LDSC resource. To assess the specific contribution of each gene set to trait heritability, we relied on the z-score of per-SNP heritability provided by S-LDSC. This z-score allows us to isolate the unique impact of each gene set while considering the influence of other annotations in the baseline model. P-values were calculated from the z-scores under the assumption of a normal distribution, and we applied the Benjamini & Hochberg procedure to control the FDR.

## DATA AVAILABILITY

Raw and processed data have been submitted to the Gene Expression Omnibus (GEO) and will be made publicly available upon publication. The code for this project is publicly available through GitHub at https://github.com/LieberInstitute/BLA_crossSpecies (Zenodo DOI: 10.5281/zenodo.13942661). Freely available web-based apps were developed to facilitate further exploration for each human (https://libd.shinyapps.io/BLA_Human), macaque (https://libd.shinyapps.io/BLA_Macaque/), and baboon (https://libd.shinyapps.io/BLA_Baboon/) datasets.

## ACKNOWLEDGEMENTS

We thank the Joint High Performance Computing Exchange (JHPCE) at Johns Hopkins University for providing computing resources for these analyses. We thank the families of Connie and Stephen Lieber and Milton and Tamar Maltz for their generous support of this work. We thank the LIBD neuropathology team, particularly James Tooke and Amy Deep-Soboslay, for curation of the brain samples and assistance with tissue dissections, and the families of the brain donors for their generosity. We thank the Tissue Distribution Program administered through Division of Comparative Medicine Pathology Services Unit at the Oregon National Primate Research Center for assistance in acquiring nonhuman primate brain tissue. We thank Jodi McBride, Allison Weiss, and Alexis Cooper for help in piloting tissue dissections. We thank the Gordon Research Conference on Amygdala Function in Emotion, Cognition, and Disease for providing a forum that ignited this collaboration. Schematic illustrations were generated using Biorender, Adobe Illustrator, and Affinity Designer.

## AUTHOR CONTRIBUTIONS

Conceptualization: VDC, RCJ, MST, SCH, KM

Methodology: RCJ, SVB, MS, KH, MRV, LBA, HT

Software: RAM, MST Validation: SVB, IDR, SCP

Formal analysis: MST, ES, VDC, RCJ

Investigation: MST, ES, VDC, RCJ, AS, LBA, SVB, MRV, MS, KH, HT

Resources: JEK, TMH, VDC

Data curation: ES, MST, RAM, AS, MT

Writing-original draft: VDC, MST, KM, SVB

Writing-review and editing: VDC, MST, KM, SVB, SCH, KM, RCJ, AS

Visualization: MST, SVB, IDR, ES

Supervision: VDC, KM, SCH

Project administration: VDC, KM, SCH

Funding acquisition: VDC, KM, SCH

