## Supplemental Figures and Tables for "Transcriptomic diversity of amygdalar subdivisions across humans and nonhuman primates"

**Running Title:** Transcriptomic characterization of amygdala cell-types across primate species.

**Authors:** Michael S. Totty[1,*], Rita Cervera Juanes[3,*], Svitlana V. Bach[2,*], Lamya Ben Ameur[4], Madeline R. Valentine[2], Evan Simons[5], McKenna Romac [5,6], Hoa Trinh[4], Krystal Henderson[5], Ishbel Del Rosario[2], Madhavi Tippani[2], Ryan A. Miller[2], Joel E. Kleinman[2,7], Stephanie Cerceo Page[2], Arpiar Saunders[4], Thomas M. Hyde[2,7,8], Keri Martinowich[2,8,9,10,+], Stephanie C. Hicks[1,11,12,13+], Vincent D. Costa[5,6,14+]

### SUPPLEMENTARY FIGURES AND TABLES

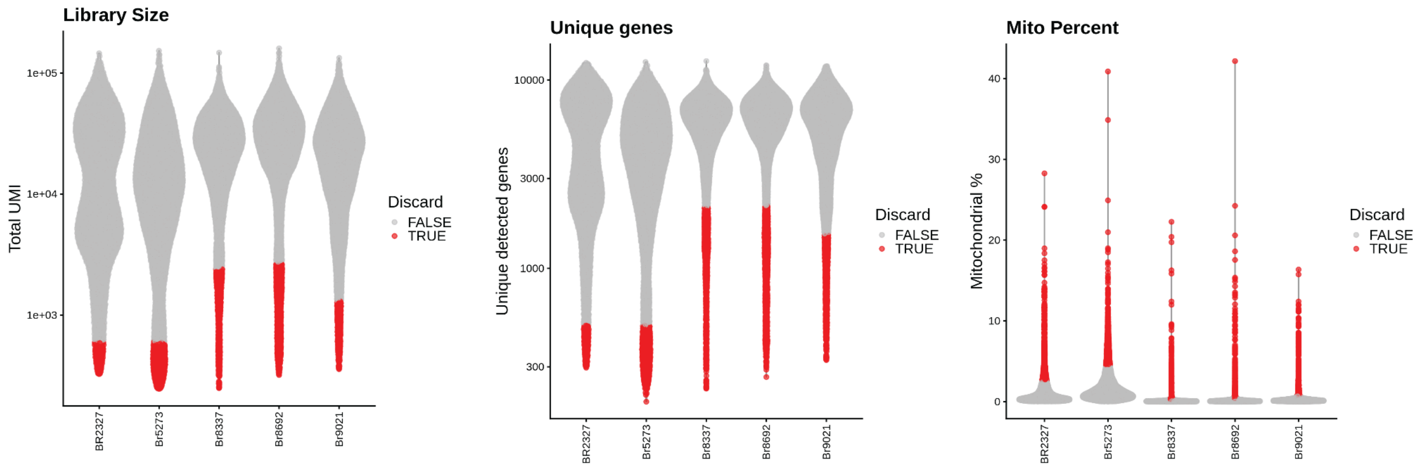


**Supplementary Figure 1: Quality control metrics for human samples.** Violin plots displaying library size, number of unique genes, and mitochondrial percent metrics across the five human samples (n=5), each from a unique donor. Red data points were considered low-quality nuclei and were excluded from downstream analyses.

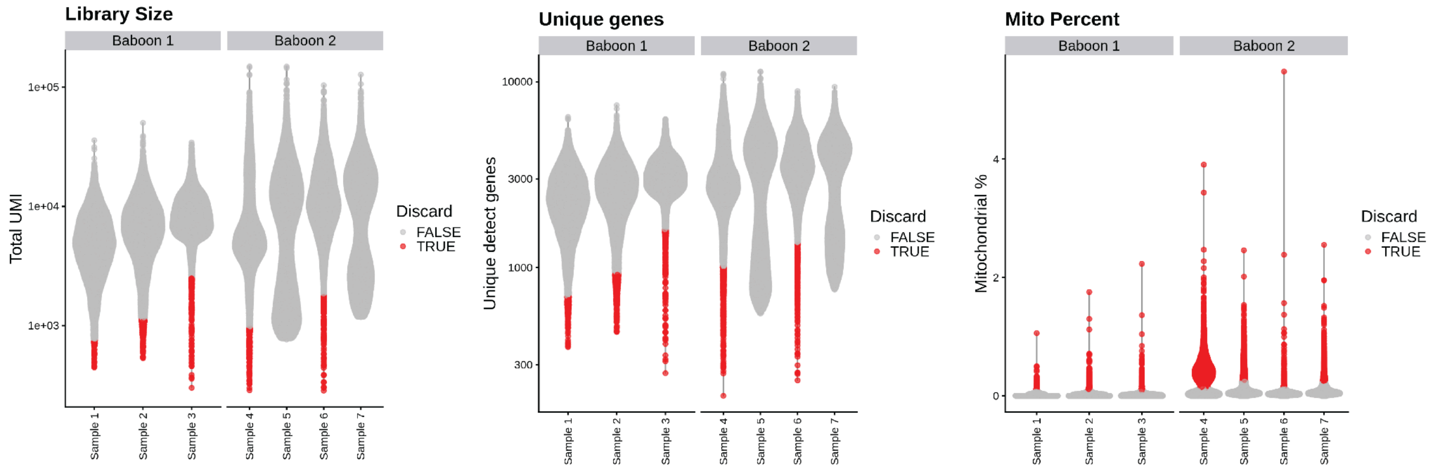


**Supplementary Figure 2: Quality control metrics for baboon samples.** Violin plots displaying library size, number of unique genes, and mitochondrial percent metrics of the seven baboon samples (n=7) across two unique donors. Red data points were considered low-quality nuclei and were excluded from downstream analyses.

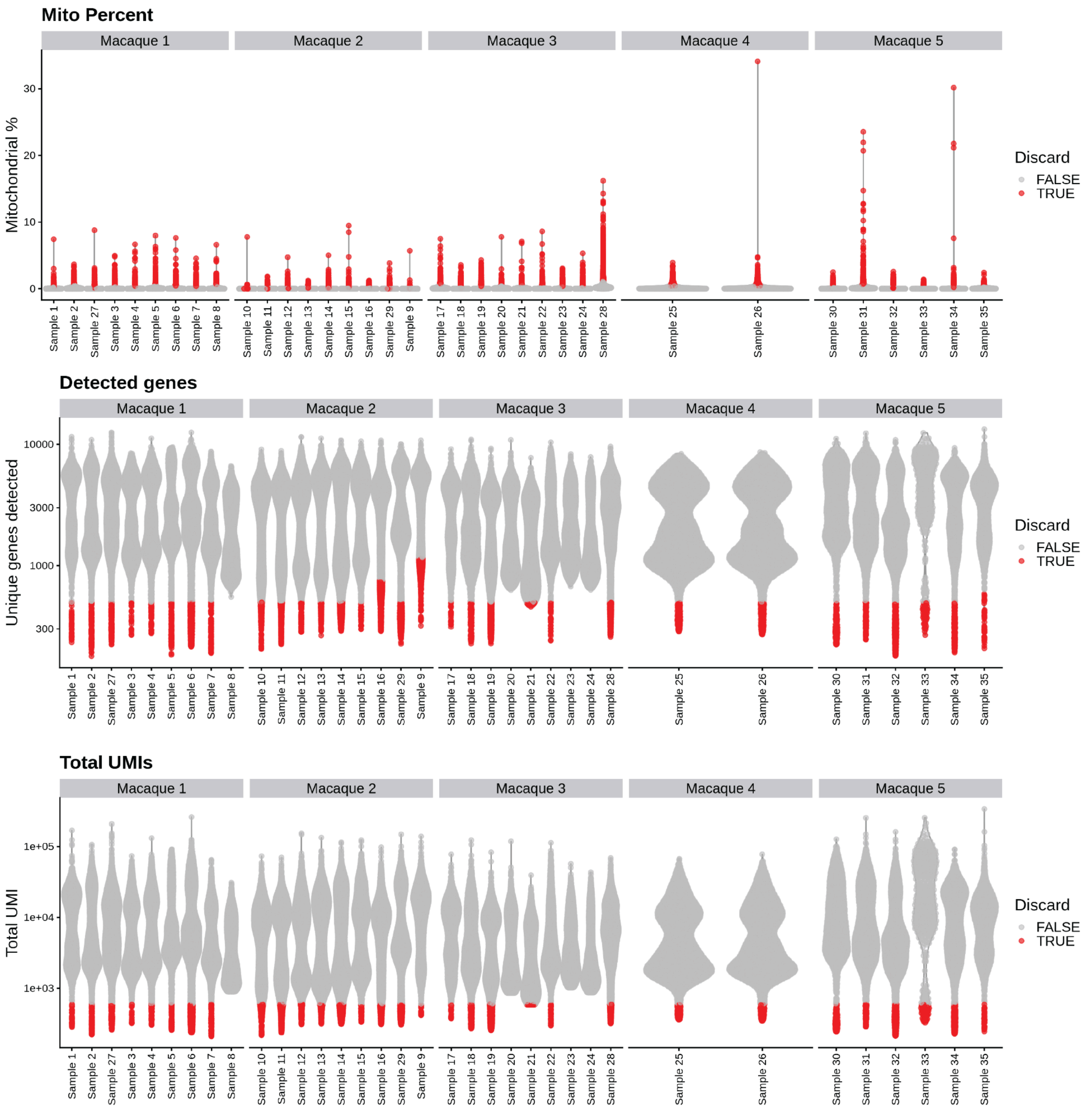


**Supplementary Figure 3: Quality control metrics for macaque samples.** Violin plots displaying library size, number of unique genes, and mitochondrial percent metrics of the thirty-five macaque samples (n=35) across five unique donors. Red data points were considered low-quality nuclei and were excluded from downstream analyses.

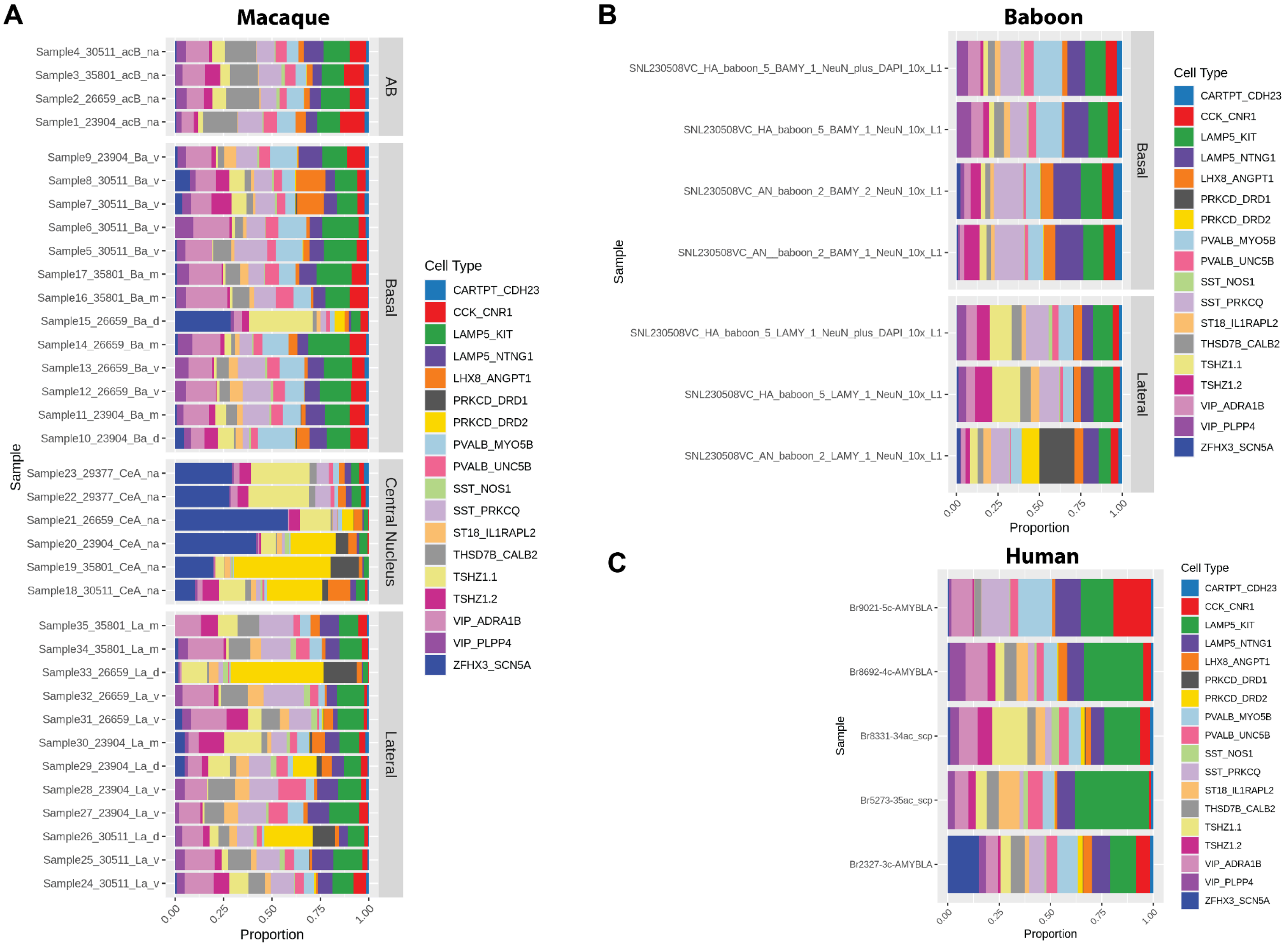


**Supplementary Figure 4: Inhibitory neuron cell type proportions across all samples.** Proportional distribution of fine inhibitory cell types across amygdala subregions in macaque, baboon, and human samples. (A) Stacked bar plots depict the proportion of each cell type within samples from different subregions of the macaque amygdala, including Accessory Basal, Basal, Central Nucleus, and Lateral punches. Sample naming scheme reflects the sample number, ONPRC ID#, subregion sampled, and if the punch was dorsal (d), intermediate (m), or ventral (v). (B) Similar analysis for baboon samples, showing cell type distribution across Basal and Lateral punches. (C) Cell type proportions in human amygdala samples which sampled the whole basolateral amygdala region. Colors represent fine cell types, as indicated in the legend, highlighting conserved patterns of cell type distribution across subregions.

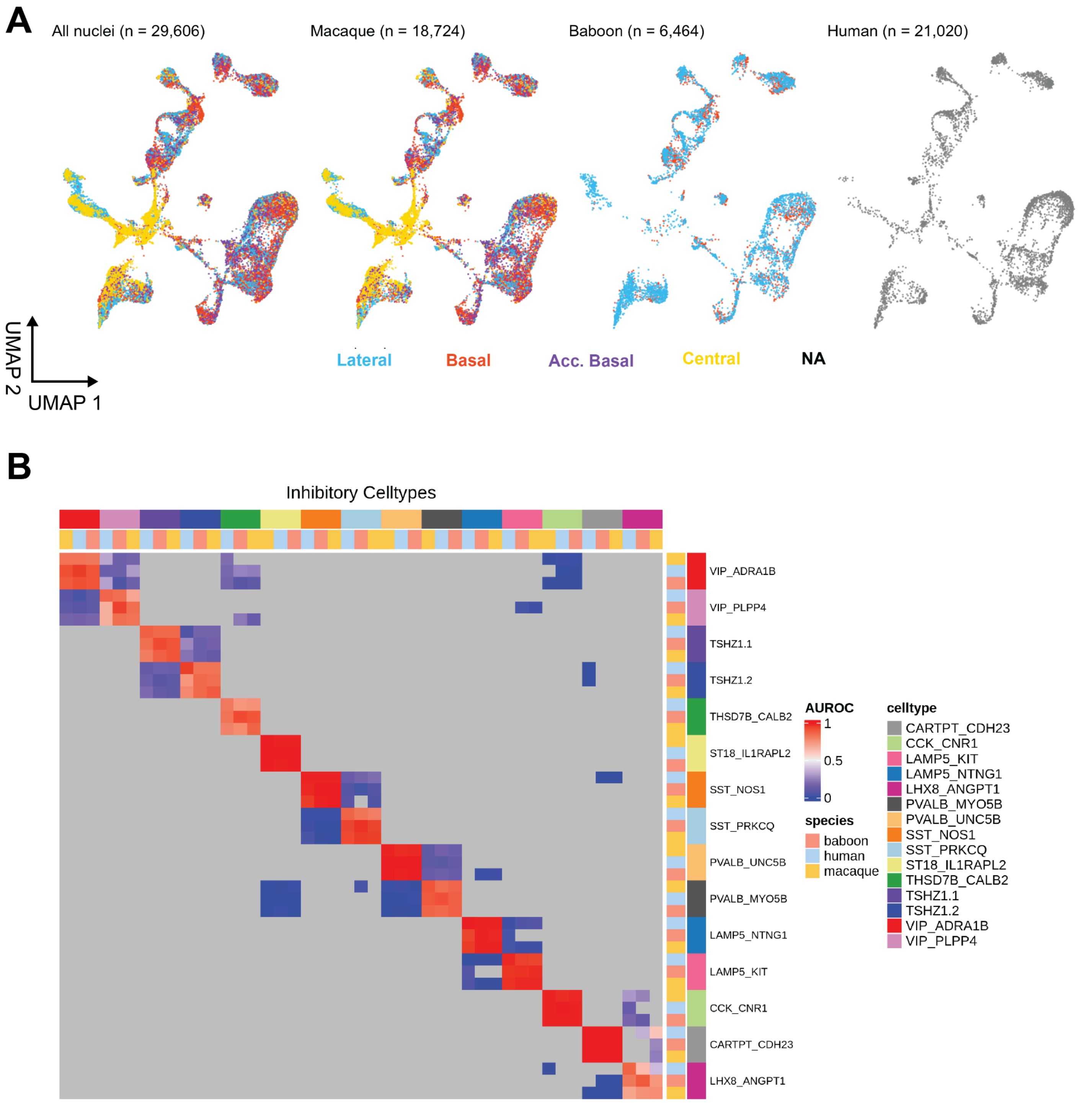


**Supplementary Figure 5: Inhibitory neurons found in the amygdala across species.** (A) UMAP visualizations of fine inhibitory cell types across species, colored by anatomical punch location. (B) Heatmap displaying one-vs-best MetaNeighbor cross-species cell type accuracy for inhibitory neurons. Each cell type was compared to the two closest matching cell types in each target dataset to test how accurately a cell type can be distinguished from its closest neighbor. Only tested cell type combinations are colored. Color scale represents the area under the receiver operating characteristic (AUROC) curve where positive (red) values indicate higher-than-chance prediction accuracy. Rows and columns indicate target and test cell types, respectively.

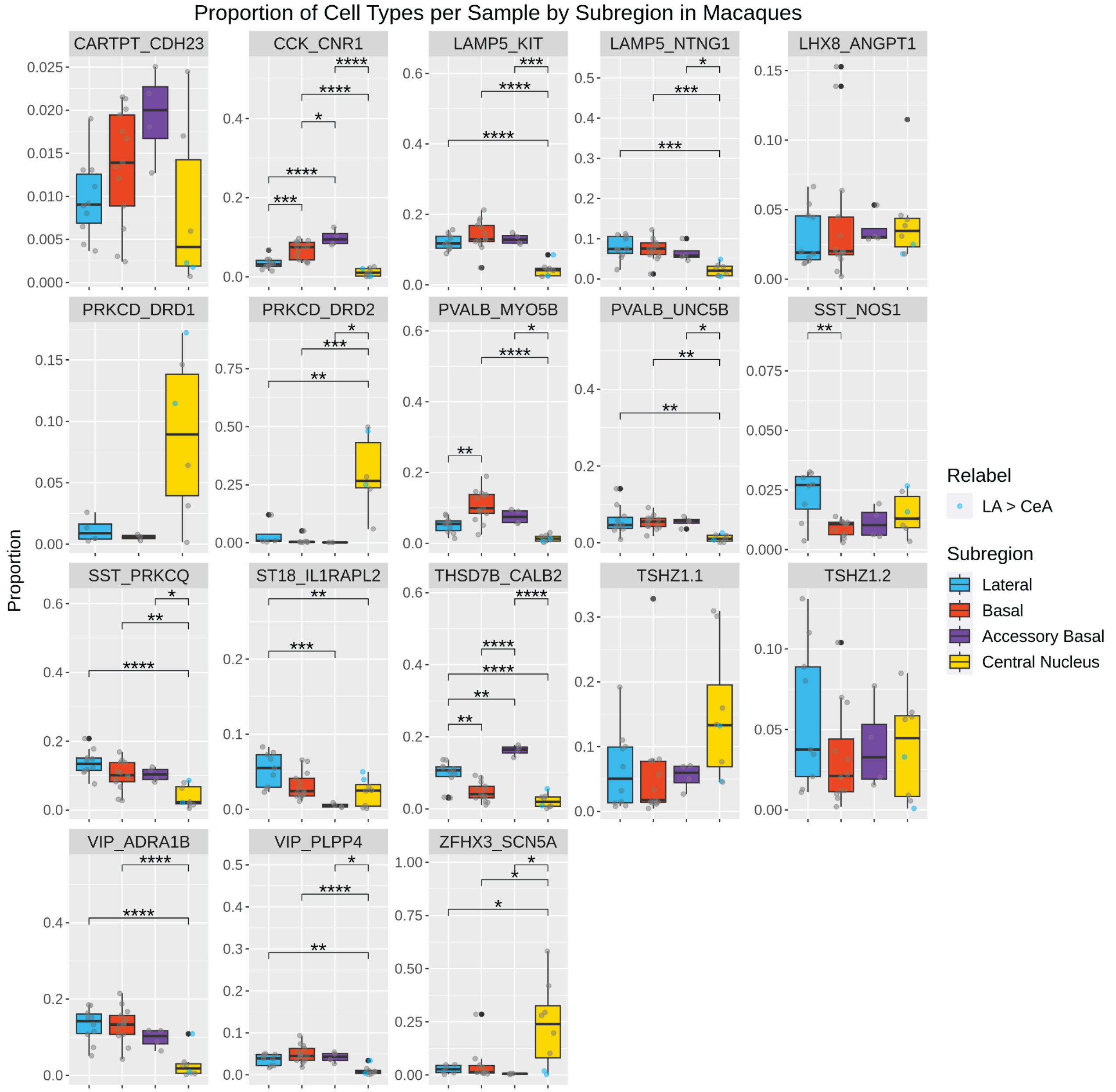


**Supplementary Figure 6: Proportion of inhibitory neurons per sample across subregion punches in Macaques.** Proportions of fine inhibitory cell types per sample across anatomical subdivisions in macaque amygdala punches. Each panel represents a different cell type, with box plots showing the proportion of nuclei per subregion: Lateral (blue), Basal (red), Accessory Basal (purple), and Central Nucleus (yellow). Grey data points represent individual samples and grey data points represent outliers. Blue data points were dorsal punches within the Lateral nucleus that were noted to have clear Central nucleus contamination. For accurate proportion estimations, these samples were relabeled as ‘Central nucleus’ only for this analysis. Significant differences between subregions are indicated by asterisks (Tukey’s HSD: * p < 0.05, ** p < 0.01, *** p < 0.001, **** p < 0.0001).

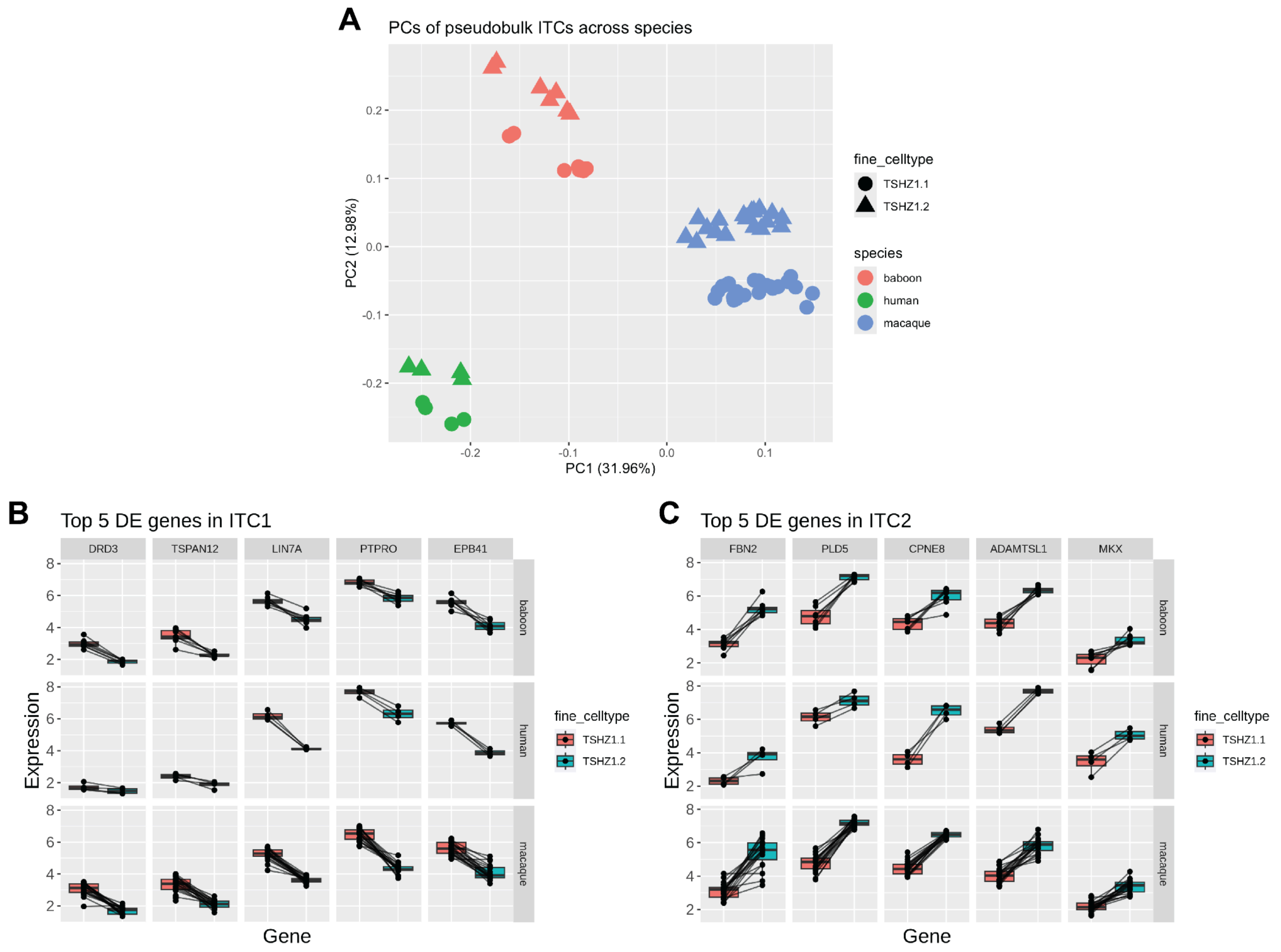


**Supplementary Figure 7: Conserved gene expression patterns among ITCs across primate species.** (A) Principal component analysis (PCA) of pseudobulk ITCs, with each point representing a sample colored by species (baboon, human, macaque) and shaped by fine cell type (TSHZ1.1 and TSHZ1.2). The first two principal components clearly separate both ITC type and species with no clear outliers, representative of high quality data. (B) Top 5 differentially expressed (DE) genes in ITC1 (TSHZ1.1), showing regularized expression levels of pseudobulk cell types across species. (C) Top 5 DE genes in ITC2 (TSHZ1.2), showing regularized expression levels of pseudobulk cell types across species. Interconnected lines represented nuclei from the same sample.

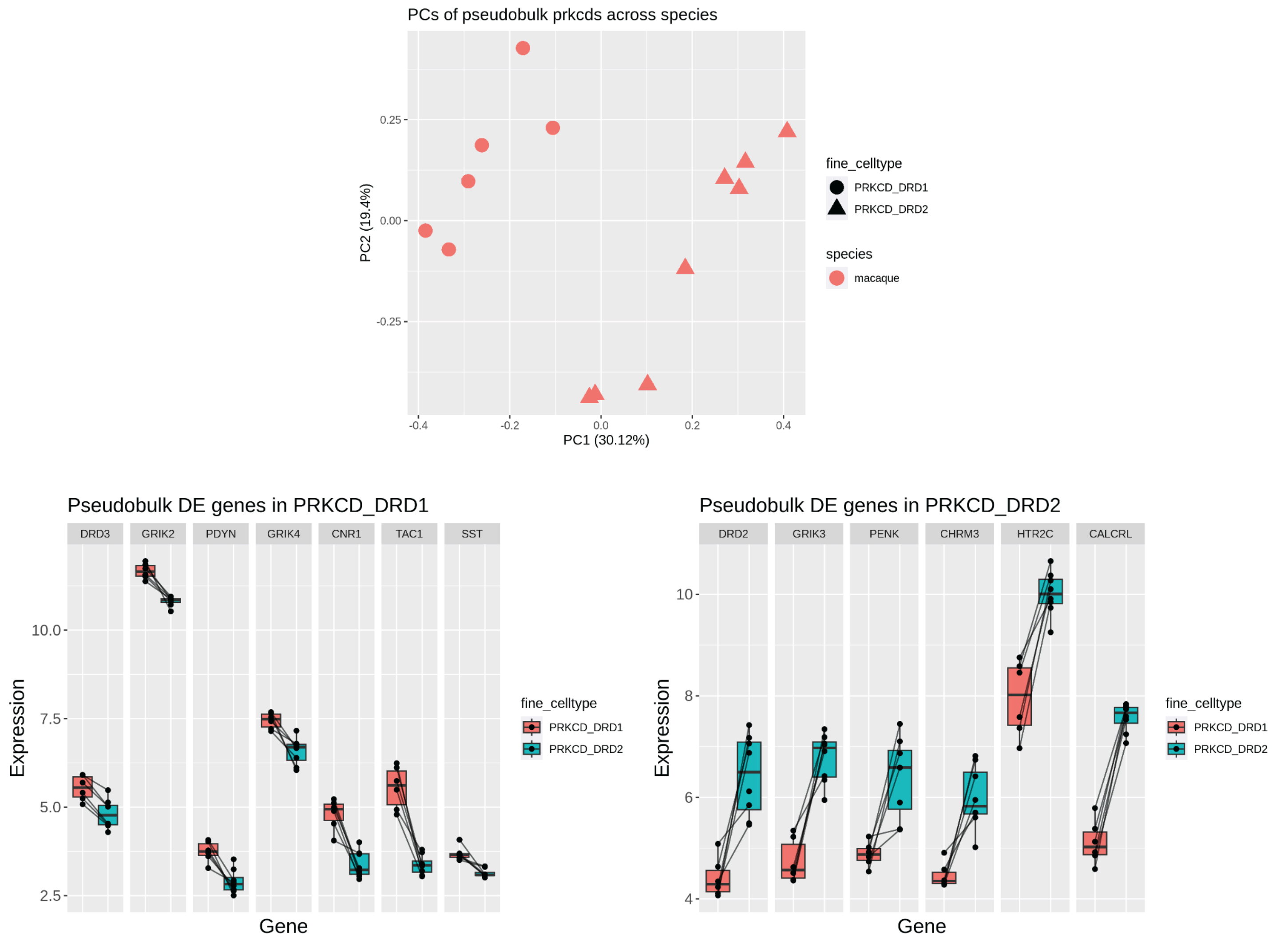


**Supplementary Figure 8: Conserved gene expression patterns among PRKCD+ inhibitory cell type primate species.** (A) Principal component analysis (PCA) of pseudobulk PRKCD+ cell types, with each point representing a sample shaped by fine cell type (PRKCD_DRD1 and PRKCD_DRD2). The first two principal components clearly separate both PRKCD+ types with no clear outliers, representative of high quality data. (B) Top 5 differentially expressed (DE) genes in PRKCD_DRD1, showing regularized expression levels of pseudobulk cell types across samples. (C) Top 5 DE genes in PRKCD_DRD2, showing regularized expression levels of pseudobulk cell types across samples. Interconnected lines represented nuclei from the same sample.

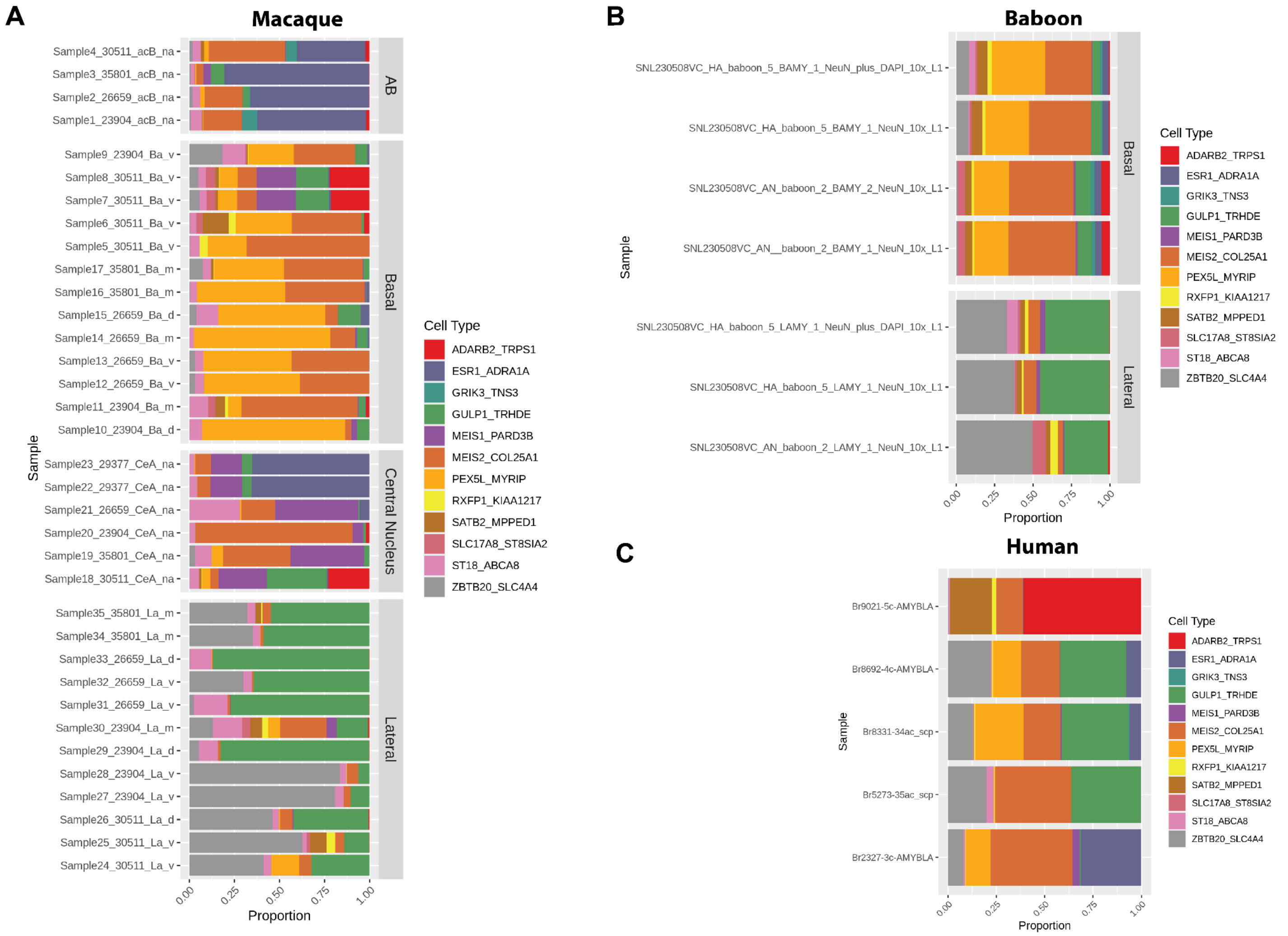


**Supplementary Figure 9: Excitatory neuron cell type proportions across all samples.** Proportional distribution of fine excitatory cell types across amygdala subregions in macaque, baboon, and human samples. (A) Stacked bar plots depict the proportion of each cell type within samples from different subregions of the macaque amygdala, including Accessory Basal, Basal, Central Nucleus, and Lateral punches. Sample naming scheme reflects the sample number, ONPRC ID#, subregion sampled, and if the punch was dorsal (d), intermediate (m), or ventral (v). (B) Similar analysis for baboon samples, showing cell type distribution across Basal and Lateral punches. (C) Cell type proportions in human amygdala samples which sampled the whole BLA. Colors represent fine cell types, as indicated in the legend, highlighting conserved patterns of cell type distribution across subregions.

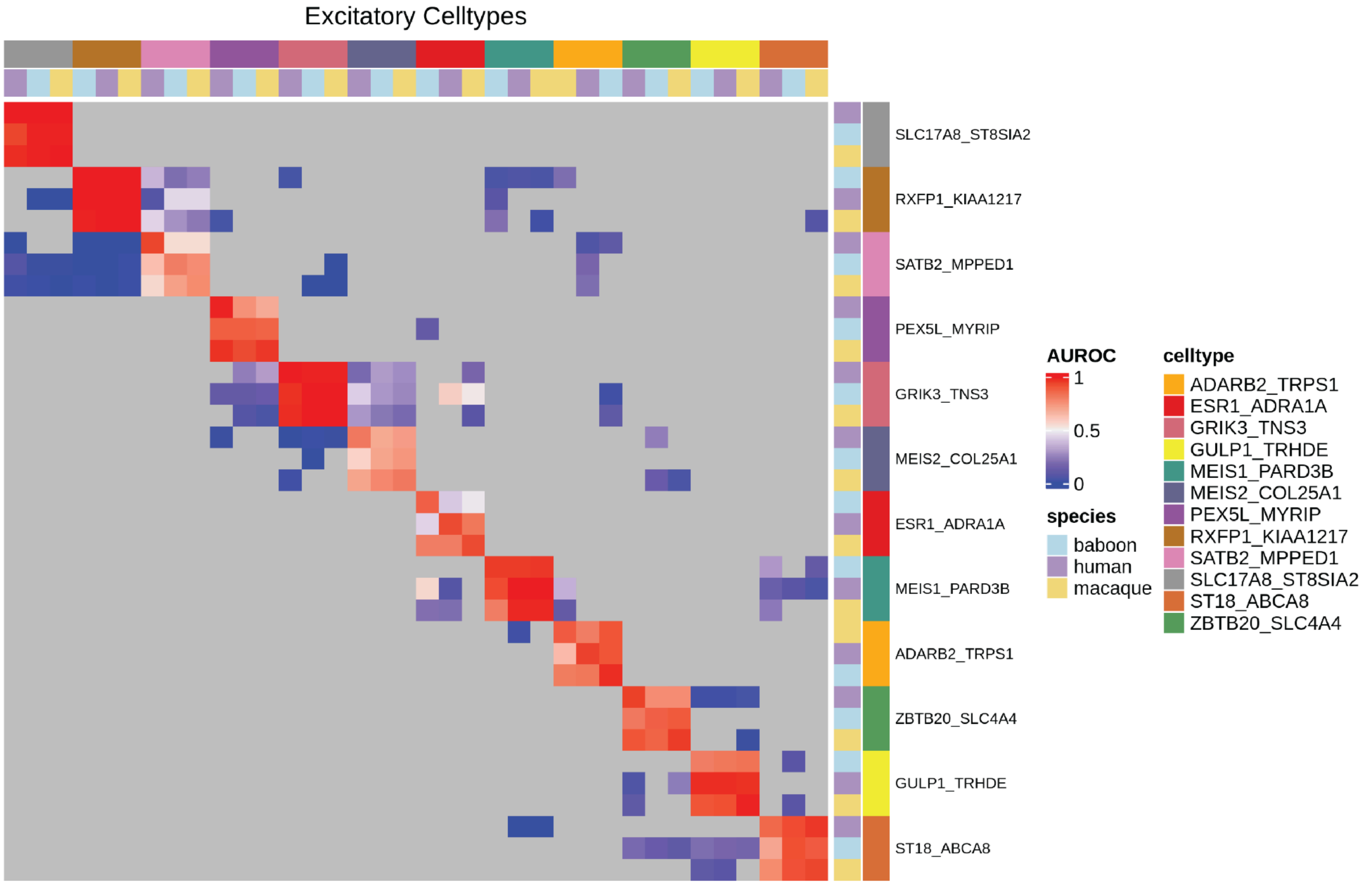


**Supplementary Figure 10: Cross-species analysis of excitatory neurons using MetaNeighbor.** (A) Heatmap displaying one-vs-best MetaNeighbor cross-species cell type accuracy for excitatory neurons. Each cell type was compared to the two closest matching cell types in each target dataset to test how easily a cell type can be distinguished from its closest neighbor. Only tested cell type combinations are colored. Color scale represents the area under the receiver operating characteristic (AUROC) curve where positive (red) values indicate higher-than-chance prediction accuracy. Rows and columns indicate target and test cell types, respectively.

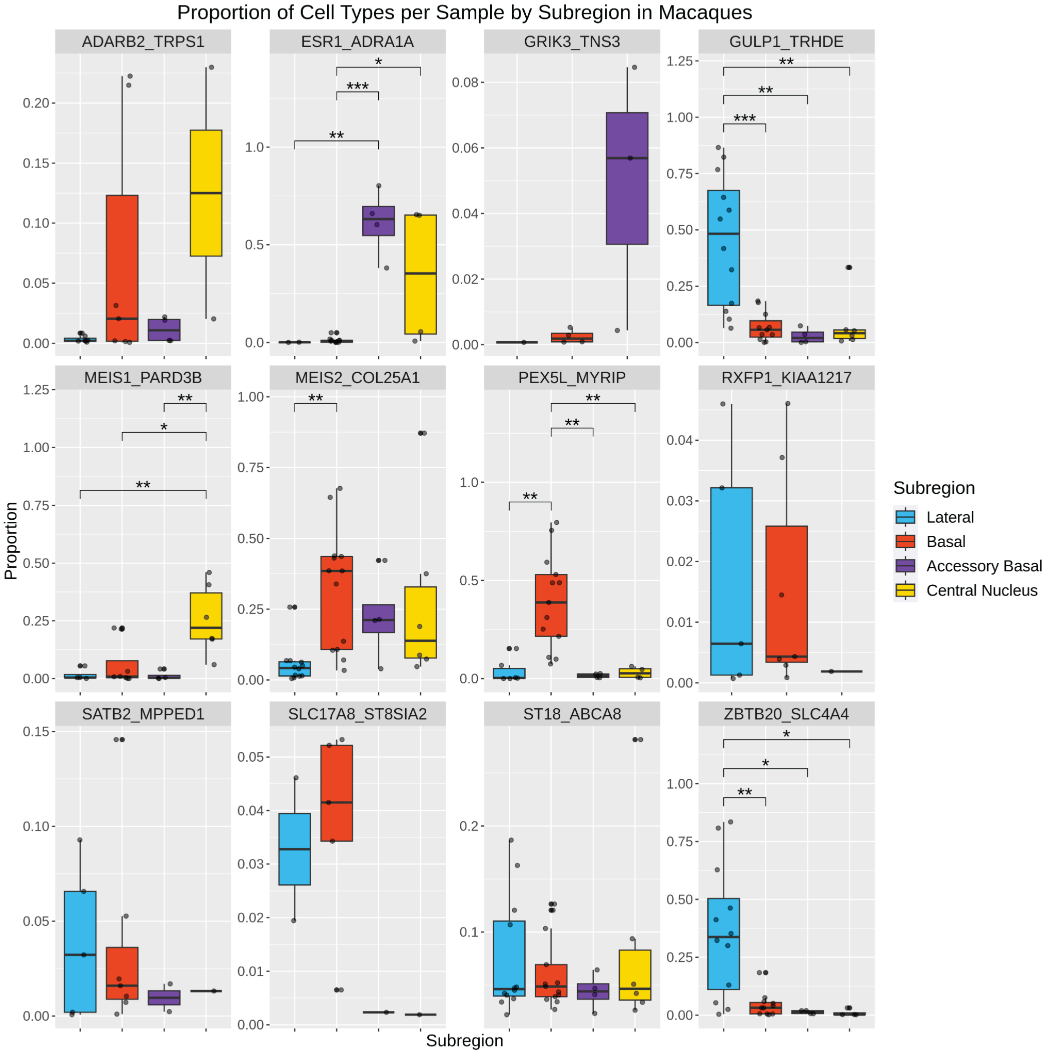


**Supplementary Figure 11: Proportion of inhibitory neurons per sample across subregion punches in Macaques.** Proportions of fine excitatory cell types per sample across anatomical subdivisions in macaque amygdala punches. Each panel represents a different cell type, with box plots showing the proportion of nuclei per subregion: Lateral (blue), Basal (red), Accessory Basal (purple), and Central Nucleus (yellow). Grey data points represent individual samples and grey data points represent outliers. Significant differences between subregions are indicated by asterisks (Tukey's HSD: * $p < 0.05$, ** $p < 0.01$, *** $p < 0.001$, **** $p < 0.0001$).

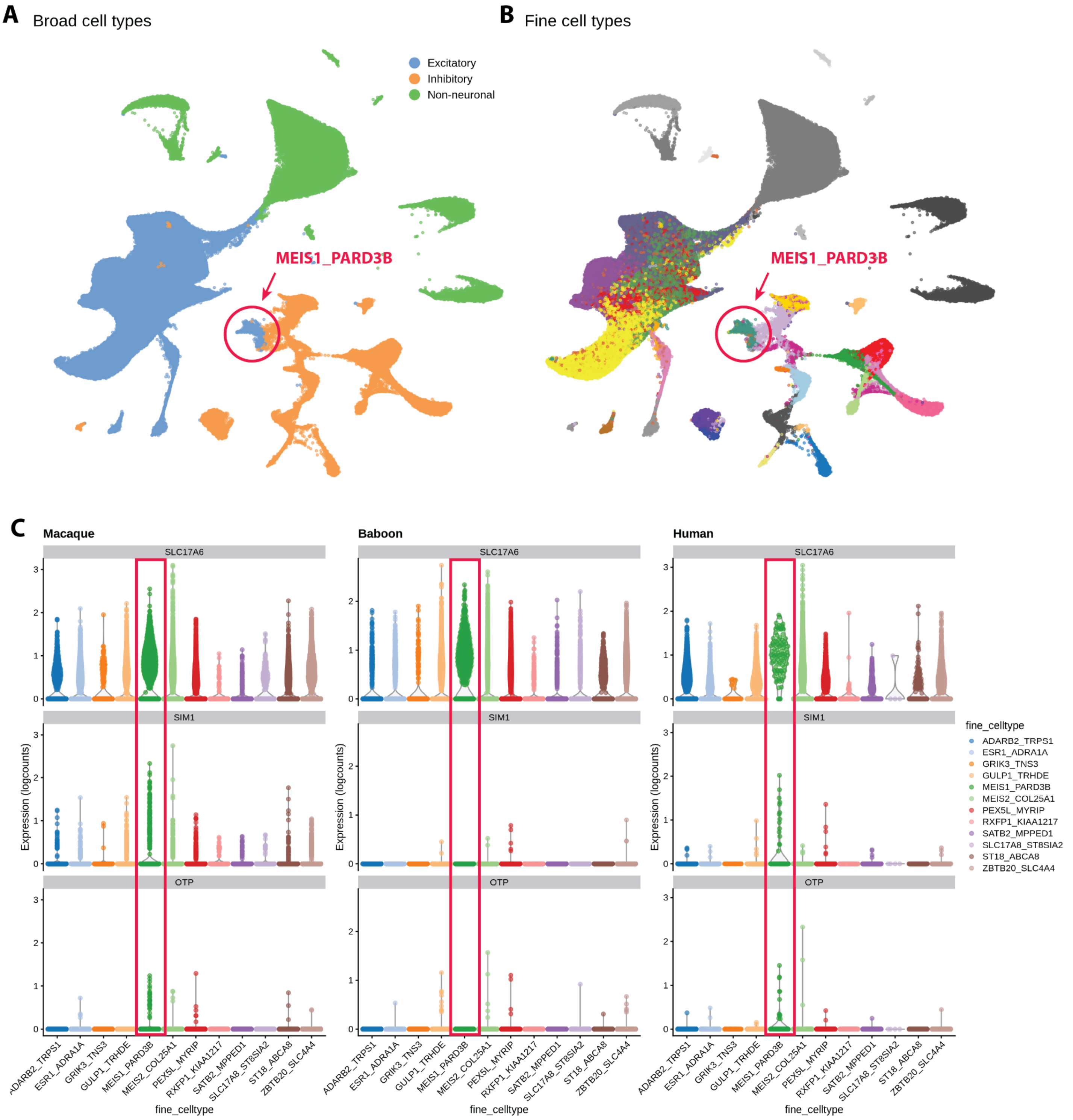


**Supplementary Figure 12: *MEIS1*⁺/*PARD3B*⁺ neurons cluster with GABAergic interneurons yet express the excitatory marker SLC17A6**. (A) UMAP visualization of broad cell types (excitatory, inhibitory, non-neuronal) and (B) fine cell types highlighting the cluster containing *MEIS1+/PARD3B+* neurons (circled and labeled in red). (C) Violin plots showing the expression of marker genes *SLC17A6*, *SIM1*, and *OTP* across different fine cell types in macaque, baboon, and human brains. The red box highlighting increased expression levels of these genes within the *MEIS1+/PARD3B+* neuron cluster, suggesting these cells are likely a class of previously characterized *SIM1*$^{+}$*/SLC17A6*$^{+}$*/GAD1*$^{-}$*/GAD2*$^{-}$ medial nucleus neurons [17].

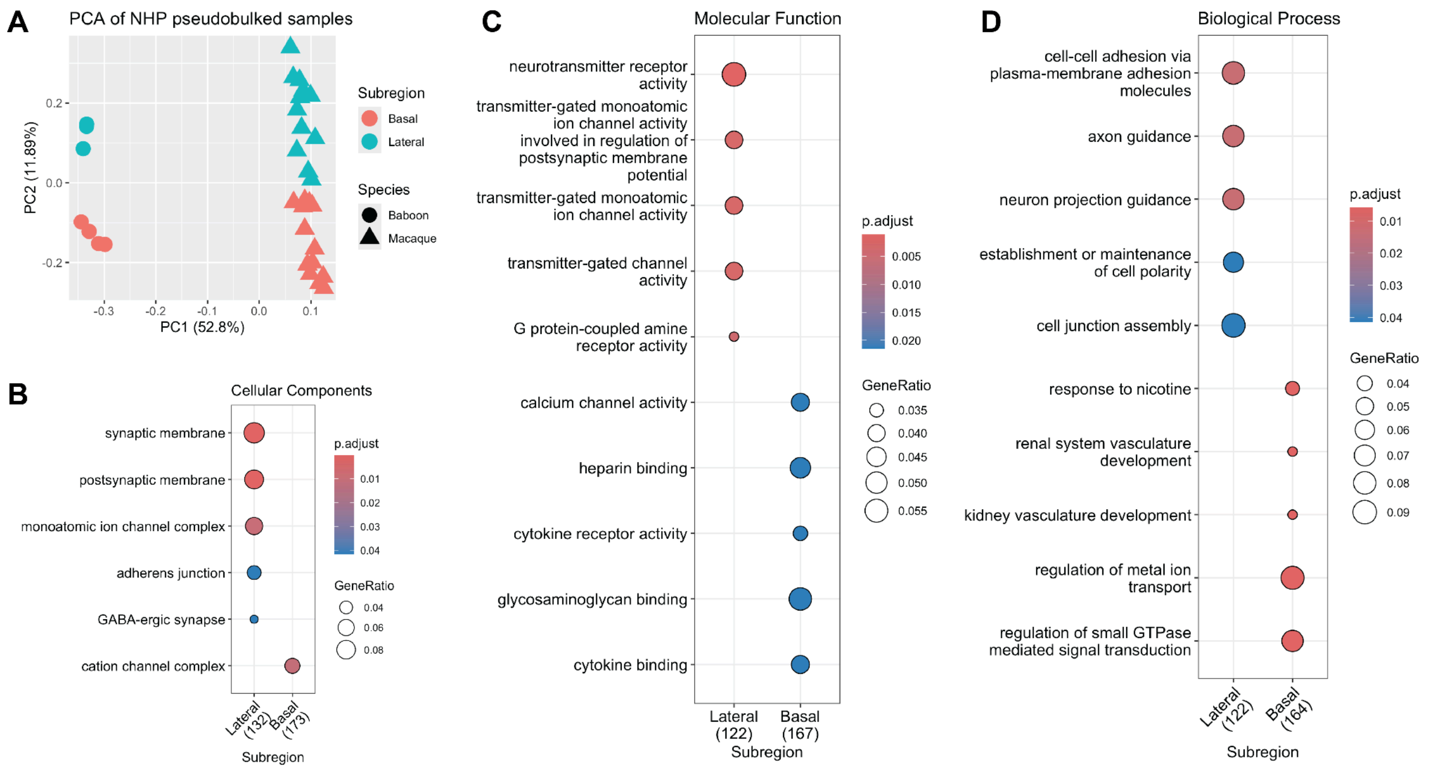


**Supplementary Figure 13: Quality control and gene ontology analysis of lateral amygdala (LA) versus basal amygdala (BA) pseudobulk data in non-human primates (NHPs).** (A) Principal component analysis (PCA) of pseudobulked excitatory neurons clearly distinguished both species and subregion with no clear outliers, indicative of high quality samples. (Basal in red and Lateral in teal; baboons represented by circles, macaques by triangles). (B) Gene Ontology (GO) terms associated with cellular components indicate significant enrichment in synaptic and membrane-related structures in either the LA or BA. (C) GO terms for molecular function highlight differences in receptor activity and channel function between subregions. (D) GO terms for biological processes illustrate distinctions in cell adhesion, axon guidance, and synapse assembly between LA and BA. Dot sizes represent gene ratios, and color gradients reflect adjusted p-values, with more significant terms in red.

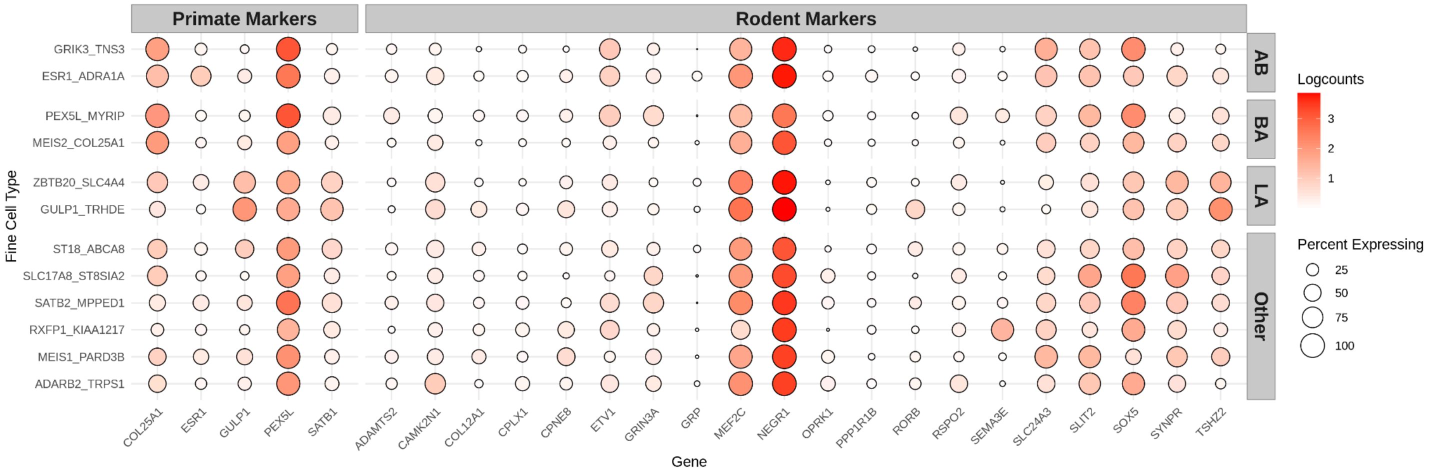


**Supplementary Figure 14: Expression patterns of primate and rodent marker genes across amygdala subregions in primates.** Dot plot visualizes the expression of marker genes across fine cell types, separated into primate (left) and rodent (right) markers. Each row represents a different cell type, and columns indicate marker genes. Dot color intensity (normalized logcounts) represents gene expression levels, while dot size indicates the percentage of cells expressing the gene. Subregions are categorized into the putative subregion cell types accessory basal (AB), basal (BA), lateral (LA), or other.

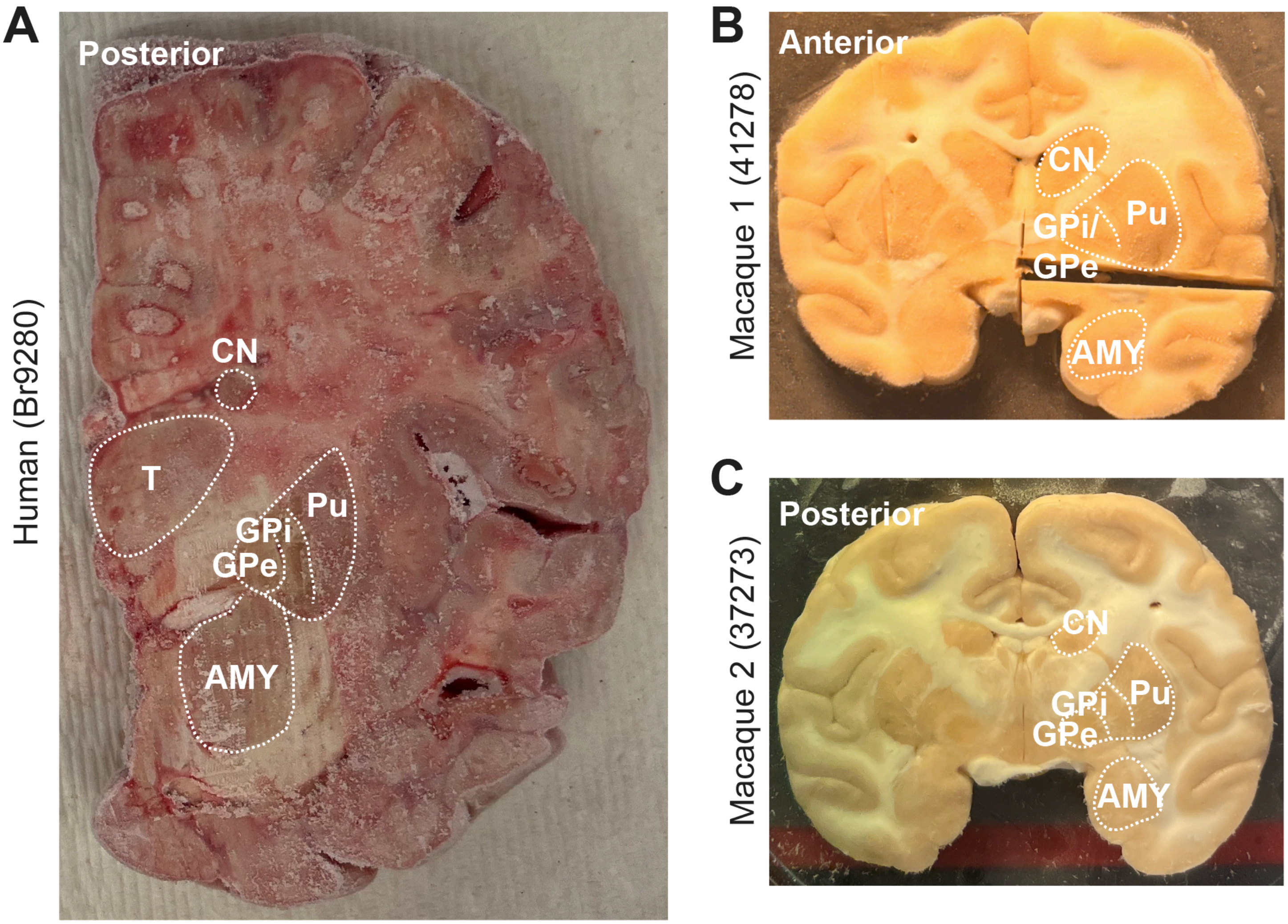


**Supplementary Figure 15. Human and macaques fresh frozen coronal brain slabs used for smFISH validation. (A)** Left human coronal brain hemisphere for donor Br9280 with indicated anatomical landmarks at the posterior level of the amygdala. **(B,C)** Coronal brain slabs for macaque 1 (41278) and macaque 2 (37273) with indicated anatomical landmarks at the anterior and posterior levels of the amygdala, respectively. AMY – amygdala, CN – caudate nucleus, GPe – globus pallidus external, GPi – globus pallidus internal, Pu – putamen, T – thalamus.

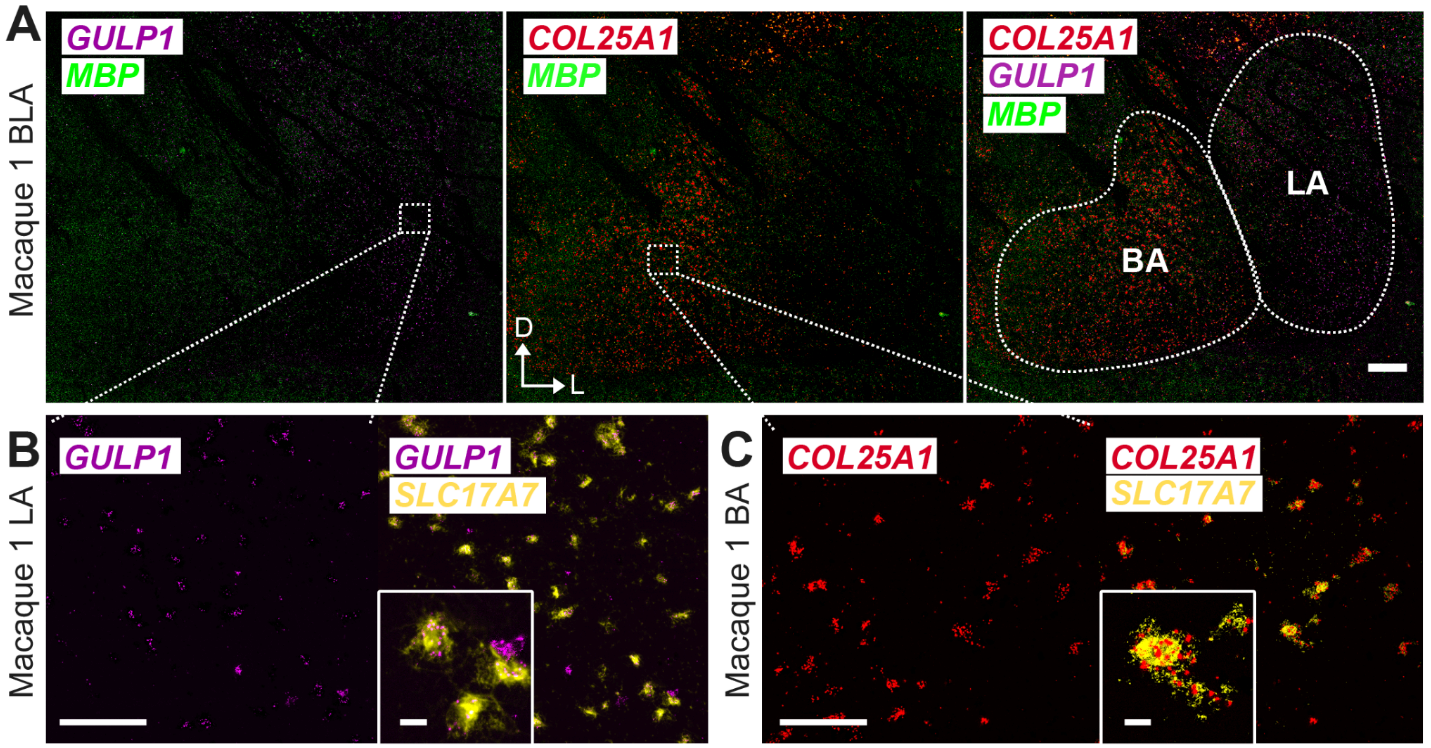


**Supplementary Figure 16. *GULP1* marker gene expression for lateral amygdala (LA) and *COL25A1* marker gene expression for basal amygdala (BA) in macaque 1. (A)** 2X smFISH images of the BLA region of the macaque brain illustrating expression of *GULP1* (magenta) in LA and *COL25A1* (red) in BA. *MBP* (green) represents white matter for anatomical landmarks. Dorval (D) and lateral (L) arrows are added for tissue directionality. White boxes represent approximate locations of zoomed-in images. Scale bar 1000 µm. **(B)** Zoomed in 40X smFISH images illustrating co-expression of *GULP1* (magenta) and *SLC17A7* (yellow) within the LA. Scale bar 100 µm. Inset illustrates representative neurons co-expressing *GULP1* and *SLC17A7*. Scale bar of inset 10 µm. **(C)** Zoomed in 40X smFISH images illustrating co-expression of *COL25A1* (red) and *SLC17A7* (yellow) within the BA. Scale bar 100 µm. Inset illustrates a representative neuron co-expressing *COL25A1* and *SLC17A7*. Scale bar of inset 10 µm.

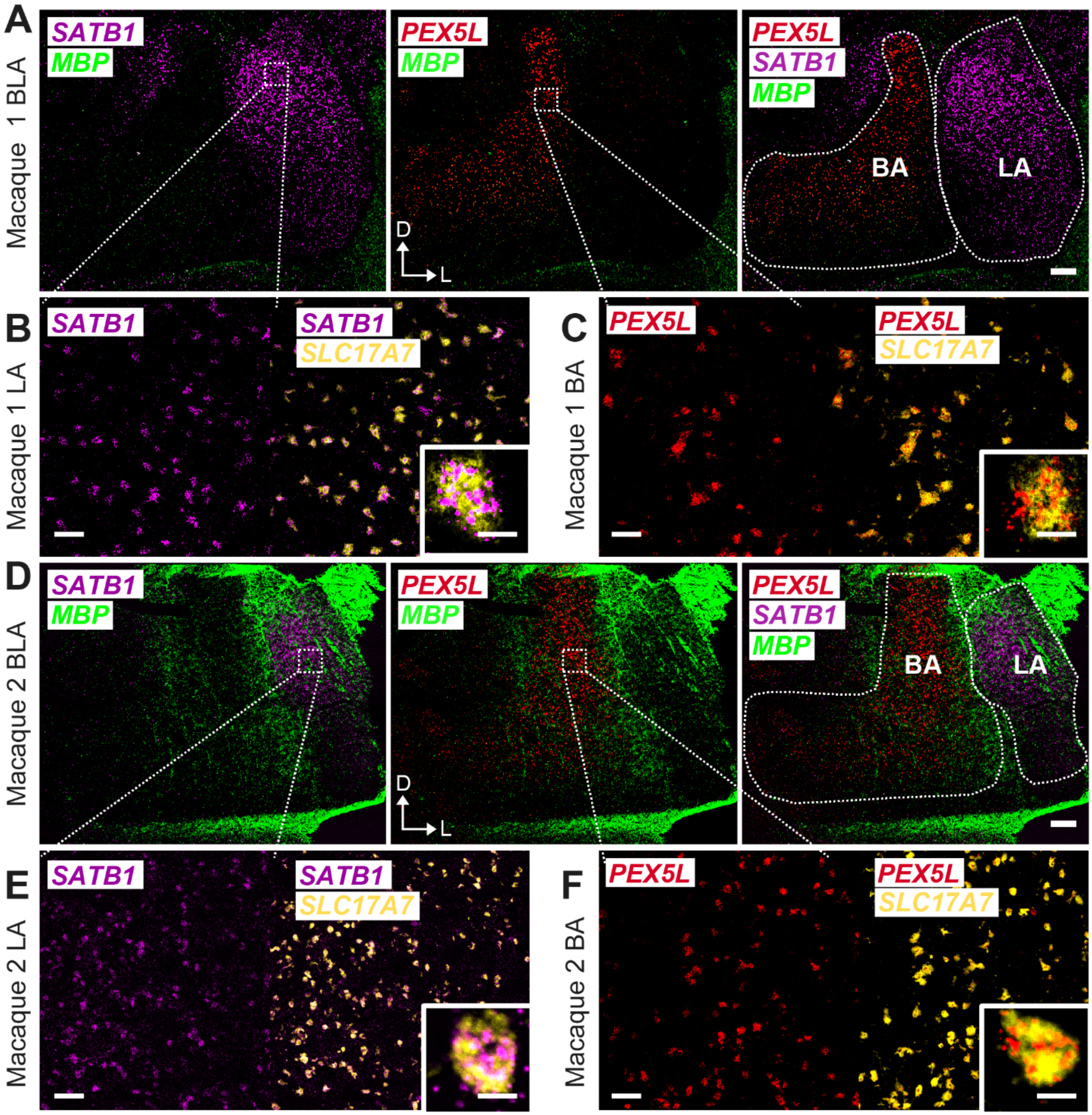


**Supplementary Figure 17. *SATB1* marker gene expression for lateral amygdala (LA) and *PEX5L* marker gene expression for basal amygdala (BA) in two macaque brains. (A, D)** 2X smFISH images of the BLA region of two macaque brains illustrating expression of *SATB1* (magenta) in LA and *PEX5L* (red) in BA. *MBP* (green) represents white matter for anatomical landmarks. Dorval (D) and lateral (L) arrows are added for tissue directionality. White boxes represent approximate locations of zoomed-in images. Scale bar 500 µm. **(B, E)** 40X smFISH images illustrating co-expression of *SATB1* (magenta) and *SLC17A7* (yellow) within the macaque LA. Scale bar 50 µm. Inset illustrates a representative neuron co-expressing *SATB1* and *SLC17A7*. Scale bar of inset 10 µm. **(C, F)** 40X smFISH images illustrating co-expression of *PEX5L* (red) and *SLC17A7* (yellow) within the macaque BA. Scale bar 50 µm. Inset illustrates a representative neuron co-expressing *PEX5L* and *SLC17A7*. Scale bar of inset 10 µm.

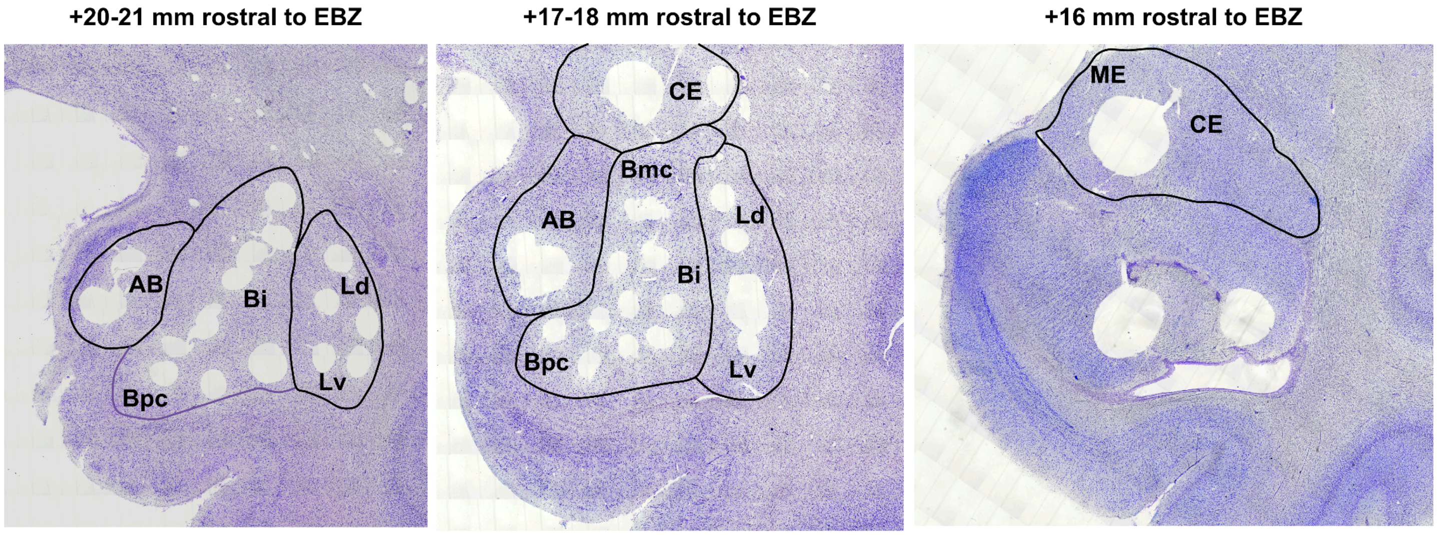


**Supplemental Figure 18. Histological reconstructions of representative tissue punches illustrating the targeted dissection strategy.** Tissue punches were taken from fresh, unfrozen tissue using gross neuroanatomical landmarks, such as the white matter tracts segregating the different nuclear subdivisions. Nissl staining after the tissue was fixed was performed to confirm accurate targeting of distinct nuclear subdivisions. Representative sections are shown from 3 different macaques, representing the rostral to caudal extent of the sampling of all four major subdivisions. Additional tissue punches from the hippocampus were taken but not sequenced. Tissue perforations that are not circled represent vasculature or tears that occurred during tissue processing. AB, accessory basal nucleus; CE, central nucleus; Bmc, magnocellular division of the basal nucleus, Bi, intermediate division of the basal nucleus; Bpc, parvocellular division of the basal nucleus; Ld, dorsal division of the lateral nucleus; Lv, ventral division of the lateral nucleus; ME, medial nucleus.

| ONPRC ID | Species | Age | Sex | Assay |
|---|---|---|---|---|
| 29377 | Macaca mulatta | 10.78 | F | snRNA-seq |
| 30511 | Macaca mulatta | 10.51 | F | snRNA-seq |
| 26659 | Macaca mulatta | 14.23 | F | snRNA-seq |
| 23904 | Macaca mulatta | 18.2 | M | snRNA-seq |
| 35801 | Macaca mulatta | 5.56 | M | snRNA-seq |
| 32273 | Macaca mulatta | 5.97 | F | RNAscope |
| 41278 | Macaca mulatta | 2.12 | M | RNAscope |
| 39944 | Papio anubis | 3.62 | F | snRNA-seq |
| 39947 | Papio anubis | 4.32 | F | snRNA-seq |

**Supplementary Table 1. Demographic information on the nonhuman primate tissue samples.** Demographic information including animal number ID in the ONPRC medical records database, species, age at time of death, sex, and assay for which the brain was utilized.

| Brain ID | Sex | Race | Age | Best RIN PFC | PMI | Assay |
|---|---|---|---|---|---|---|
| Br5273 | F | Caucasian | 53.89 | 7.0 | 34.5 | snRNA-seq |
| Br8331 | M | Hispanic | 47.5 | 7.2 | 12.5 | snRNA-seq |
| Br2723 | M | Caucasian | 50.4 | 7.1 | 25.5 | snRNA-seq |
| Br9021 | M | African American | 70.74 | 7.6 | 15.5 | snRNA-seq |
| Br8692 | M | African American | 48.21 | 8.3 | 10.5 | snRNA-seq |
| Br9280 | M | Caucasian | 66.6 | 9.3 | 25.5 | RNAscope |

**Supplementary Table 2. Human brain donor demographic information.** Demographic information on brain donors including Brain ID, sex (F, female; M, male), race (CAUC, Caucasian; HISP, Hispanic; AA, African American), age at the time of death, best RNA integrity number (RIN) measured in the prefrontal cortex (PFC), post-mortem interval (PMI), and assay for which the brain was utilized.
